# Quantifying exposure of bumblebee (*Bombus* spp.) queens to pesticide residues when hibernating in agricultural soils

**DOI:** 10.1101/2022.05.10.491216

**Authors:** Sabrina Rondeau, Nicolas Baert, Scott McArt, Nigel E. Raine

## Abstract

Exposure to pesticides is a major threat to bumblebee (*Bombus* spp.) health. In temperate regions, queens of many bumblebee species hibernate underground for several months, putting them at potentially high risk of exposure to soil contaminants. The extent to which bumblebees are exposed to residues in agricultural soils during hibernation is currently unknown, which limits our understanding of the full pesticide exposome for bumblebees throughout their lifecycle. To generate field exposure estimates for overwintering bumblebee queens to pesticide residues, we sampled soils from areas corresponding to suitable likely hibernation sites at six apple orchards and 13 diversified farms throughout Southern Ontario (Canada) in fall 2019-2020. Detectable levels of pesticides were found in 65 of 66 soil samples analysed for multi-pesticide residues (UPLC-MS/MS). A total of 53 active ingredients (AIs) were detected in soils, including 27 fungicides, 13 insecticides, and 13 herbicides. Overall, the frequency of detection, residue levels (median = 37.8 vs. 2.2 ng/g), and number of pesticides per sample (mean = 12.3 vs. 4.3 AIs) were highest for orchard soils compared to soils from diversified farms. Ninety-one percent of samples contained multiple residues, including mixtures of insecticides and fungicides that might lead to synergistic effects. Up to 29 different AIs were detected per soil sample. Our results suggest that when hibernating in agricultural areas, bumblebee queens are very likely to be exposed to a wide range of pesticide residues in soil. Our study indicates the importance of empirically testing the potential effects of pesticide residues in soils for hibernating bumblebee queens, using field exposure data such as those generated here. The differences in exposure that we detected between cropping systems can also be used to better inform regulations that govern the use of agricultural pesticides, notably in apple orchards.

## INTRODUCTION

Many bumblebee, *Bombus* spp. (Hymenoptera: Apidae), species within North America and Europe have experienced significant population declines in recent decades (Cameron *et al*. 2020 and references therein), raising concerns over the fate of wild bee populations, agricultural production, and the maintenance of biodiversity. Widespread pesticide usage is one of the multiple interacting factors that have been linked to decreases in bumblebee populations (Blacquière *et al*. 2012; Goulson *et al*. 2015; Baron *et al*. 2017). Although the extent to which pesticide exposure contributes to these declines is not fully understood, studies have found that exposure to insecticides can compromise bumblebees’ learning and foraging capabilities, colony initiation and reproductive success, as well as their efficiency in pollinating crops (Godfray et al. 2014; Stanley et al. 2015; Siviter et al. 2021; Camp & Lehmann 2021). Exposure to fungicides, a class of pesticides generally considered safer for bees, can also have detrimental effects on bumblebees by impairing colony performance and increasing their susceptibility to other pesticides and gut parasite infections (Belsky & Joshi 2020). Even ‘inactive’ ingredients, used as co-formulants in some of the most widely used herbicide products, have been reported to cause high levels of bumblebee mortality (Straw et al. 2021).

In the agricultural environment, bumblebees can be exposed to pesticides either orally, through the consumption of contaminated pollen and nectar, or via contact with plant surface residues, foliar sprays, or dust particles generated from the planting of treated seeds (Godfray et al. 2014; Gradish *et al*. 2019). Bumblebees commonly forage on crops treated with various pesticides (David *et al*. 2016; Baron *et al*. 2017; Ellis *et al*. 2017) and even minimal agricultural activity in conservation areas have been reported to expose bumblebees to a range of insecticides, fungicides, and herbicides (Main *et al*. 2020a). Pesticide residues have been detected in the food stores of bumblebee colonies placed next to agricultural crops (David *et al*. 2016; Thompson *et al*. 2016; Ellis *et al*. 2017), in puddles of water near agricultural fields (Samson-Robert *et al*. 2014, McCune *et al*. 2021) from which bumblebees may drink, as well as in the tissues of wild and commercial bumblebees on arable farms (David *et al*. 2015, 2016; Botias *et al*. 2017; Main *et al*. 2020b). In underground-hibernating species, bumblebee queens may also be exposed to residues in soil during hibernation (Gradish *et al*. 2019), but this route of exposure remains largely unexplored and the extent to which bumblebees are exposed to residues in soil has yet to be documented. This is an important research gap as considering the full pesticide exposome for bumblebees throughout their lifecycle, including exposure through soil during hibernation, is critical to estimate the risk pesticides represent for bumblebees and develop proper management strategies. Understanding risk for bumblebee queens is particularly important considering that the success of a colony depends entirely on the queen’s survival and ability to initiate a colony the following season.

In temperate regions, bumblebees have an annual lifecycle that includes a solitary phase during which only the young queens produced at the end of summer survive the winter by hibernating underground for six to nine months (Plath 1927; Alford 1969). In early spring, these queens emerge from hibernation to initiate new colonies. Therefore, bumblebee queens spend most of their lifecycle underground during hibernation, putting them at potentially high risk of exposure to soil contaminants (Gradish *et al*. 2019). Considering the physiological stress of the extended hibernation period for bumblebee queens, any effect of pesticides is likely to be amplified under such conditions. Exposure to soil residues is also probable for bumblebee larvae and adults of bumblebee species that nest underground, but the likelihood of such exposure is deemed comparatively low (Gradish *et al*. 2019).

Within agricultural landscapes, bumblebees primarily nest and hibernate in uncropped field margins or surrounding patches of woodlands, meadows, gardens, or pastures (Liczner & Colla 2019; Purvis *et al*. 2020) and are unlikely to hibernate directly within crop fields. This makes estimating the exposure of queens to pesticides in soil from the current literature impractical considering that existing residue data most often reflect residues within field soils (Gradish *et al*. 2019). A better approach to estimate exposure would be to quantify residue levels in soils sampled from suitable hibernation sites for bumblebee queens in agricultural settings.

Although characteristics of overwintering sites have only been described for a few North American and European species (Liczner & Colla 2019), many similarities exist across studies. Bumblebee queens have most often been observed hibernating in North-facing sloping sites (e.g., banks, walls of ditches, mounds of soil) and, occasionally, in slopes facing east or west (Sladen 1912; Plath 1927; Alford 1969). Preferences regarding the height and inclination of slopes likely vary among bumblebee species, with some species observed hibernating almost exclusively in steep banks and others in more gently sloping ground (Alford 1969). Queens have been found hibernating in sandy, well-drained, and loose soils from sites often shaded from direct sunlight, usually by nearby trees, and from sites that are free of dense ground vegetation (Sladen 1912; Plath 1927; Alford 1969). Although the depth of the hibernation chamber (hibernaculum) may vary according to ground conditions, regions, and species (Liczner & Colla 2019), bumblebee queens have been found hibernating at depths between 2 and 15 cm, and most frequently less than 10 cm (Plath 1927; Alford 1969).

In Southern Ontario, agricultural field margins appear to provide bumblebees with food, nesting, and hibernating resources comparable to those of semi-natural habitats (Purvis et al. 2020). Bumblebees are common visitors of apples, berries, squash, and many other crops and, as such, are likely to hibernate near farms growing these crops (Blaauw & Isaacs 2014; Gervais 2021; Willis Chan & Raine 2021). Here, we quantified pesticide residues in agricultural soils to understand exposure for hibernating bumblebee queens and compared these residues between apple orchards and diversified farms, two cropping systems commonly encountered in Ontario and differing in their pesticide use patterns. Considering that pesticide use is often more intensive in permanent crops (Farm and Food Care Ontario 2015; OECD 2019), we hypothesized that pesticide content will be higher in orchard soils.

## MATERIALS AND METHODS

### Sites and soil sampling

We sampled soil from 19 conventional farms located in Southern Ontario, Canada (Fig. S1) between October 7 and 9, 2019 and between September 22 and October 9, 2020. These included six apple orchards and 13 farms growing multiple crops, such as corn (*Zea mays*), soybeans (*Glycine max*), strawberries (*Fragaria* spp.), pumpkins and squash (*Cucurbita* spp.), and other vegetables.

At each farm, we collected soil adjacent to agricultural fields from at least three different areas corresponding to suitable likely bumblebee queen hibernation sites, using a list of criteria based on available literature (see references in introduction). We searched for sloping ground with a minimum inclination of 10° from the horizontal and a slope that was facing north (preferably), west, or east. The chosen sites were shaded from direct sunlight, most often by trees, and were free of dense living ground vegetation. The soil was well-drained (no clear accumulation of water on soil surface or excessive soil moisture) and rather loose (i.e., easy to sample with our manual core sampler without too much resistance). At each sampling site (n=66), we collected approximately 600 ml of soil (0-10 cm) in plastic bags, using a soil corer. We also recorded data on the slope inclination of the site and the distance from the edge of the nearest cultivated field (or apple rows). All samples were kept on ice until their arrival at the laboratory.

### Observations of queens searching for hibernation sites

In 2020, field observations were made to find and characterize real bumblebee hibernation sites. At each of the eight farms sampled in 2020, we walked along the field edges and the margins of forested areas and other uncropped areas between 10:00 and 15:00 on days when weather conditions were suitable (sunny or partly cloudy with minimal wind). We walked these sites at a steady pace for a total of six hours per farm (divided between two or three observers) in search of bumblebee queens that were digging in the ground. Although many bumblebees (workers, males, and sometimes queens) were observed at each farm, we only observed one queen (*Bombus impatiens*) digging in soil at one site. Data on site characteristics were collected and soil was sampled as described above.

### Sample preparation and analyses

At the end of each sampling day, all soil samples were homogenized and divided into subsamples. Soil pH and humidity were measured following standard methods (Michigan State University 2019a,b) and the rest of the soil was frozen at −20°C until analysis. Within three months from sampling, soil subsamples (50 g) were shipped on dry ice to the Chemical Ecology Core Facility at Cornell University (Ithaca, NY, USA) for multi-pesticide analysis. Frozen soil samples were extracted using the EN 15662 QuEChERS procedure (European Committee for Standardization 2018) and screened for 247 pesticides and metabolites (Table S1) by liquid chromatography – tandem mass spectrometry (UHPLC-MS/MS) (see Appendix S1 for details). Subsamples (300 g) were also sent to A&L Canada Laboratories (London, ON, Canada) where soil texture (%sand, %silt, and %clay) was characterized using the standard hydrometer method and the organic matter (OM) content measured by loss on ignition.

### Data analysis

All pesticide residues above the respective limits of detection (LODs) were considered in data analysis; data entries above LODs but below limits of quantification (LOQs) were replaced by their respective LOQ values. Statistical analyses were conducted using the R software (R Core Team 2021). To compare soil contamination between cropping systems, we ran generalized linear mixed models (GLMMs) with the total pesticide content (sum of all residue concentrations) and the number of active ingredients (AIs) detected per sample as response variables, using either a gamma or a poisson distribution, as appropriate. Models were fitted using the type of cropping system and distance to the nearest crops as fixed effects and the farm as a random effect.

We performed Principal Component Analyses (PCAs) and Spearman’s rank correlation analyses to explore the potential relationships between soil pesticide content and soil (pH, humidity, % silt and clay, OM content), site (slope inclination, distance to nearest field), and pesticide properties. Pesticide properties (Table S2) were obtained from the Pesticide Properties Database (Lewis *et al*. 2016) and other relevant literature and included: half-life time in soil (DT50, days, indicator of persistence in soil), solubility in water at 20°C (Sw, mg/L), octanol-water partition coefficient at pH 7 and 20°C (LogP, measure of lipophilicity and indicator of bioaccumulation potential), vapour pressure at 20°C (Vp, mPa, indicator of volatility), GUS index (indicator of leaching potential), and organic carbon-water partition coefficient (Koc, ml/g, indicator of soil adsorption and mobility). Of all 53 AIs detected in soil samples, one (Griseofulvin) was excluded from the analysis due to missing values of some properties.

## RESULTS

### Pesticide content in soils

Of the 66 soil samples collected, all but one contained detectable residues of at least one AI (Fig. 1), with 90.9% (60/66) of the samples containing multiple residues. Overall, herbicides were the most frequently detected (93.9% of samples), followed by insecticides (81.8%), and fungicides (59.1%).

**Figure 1.**
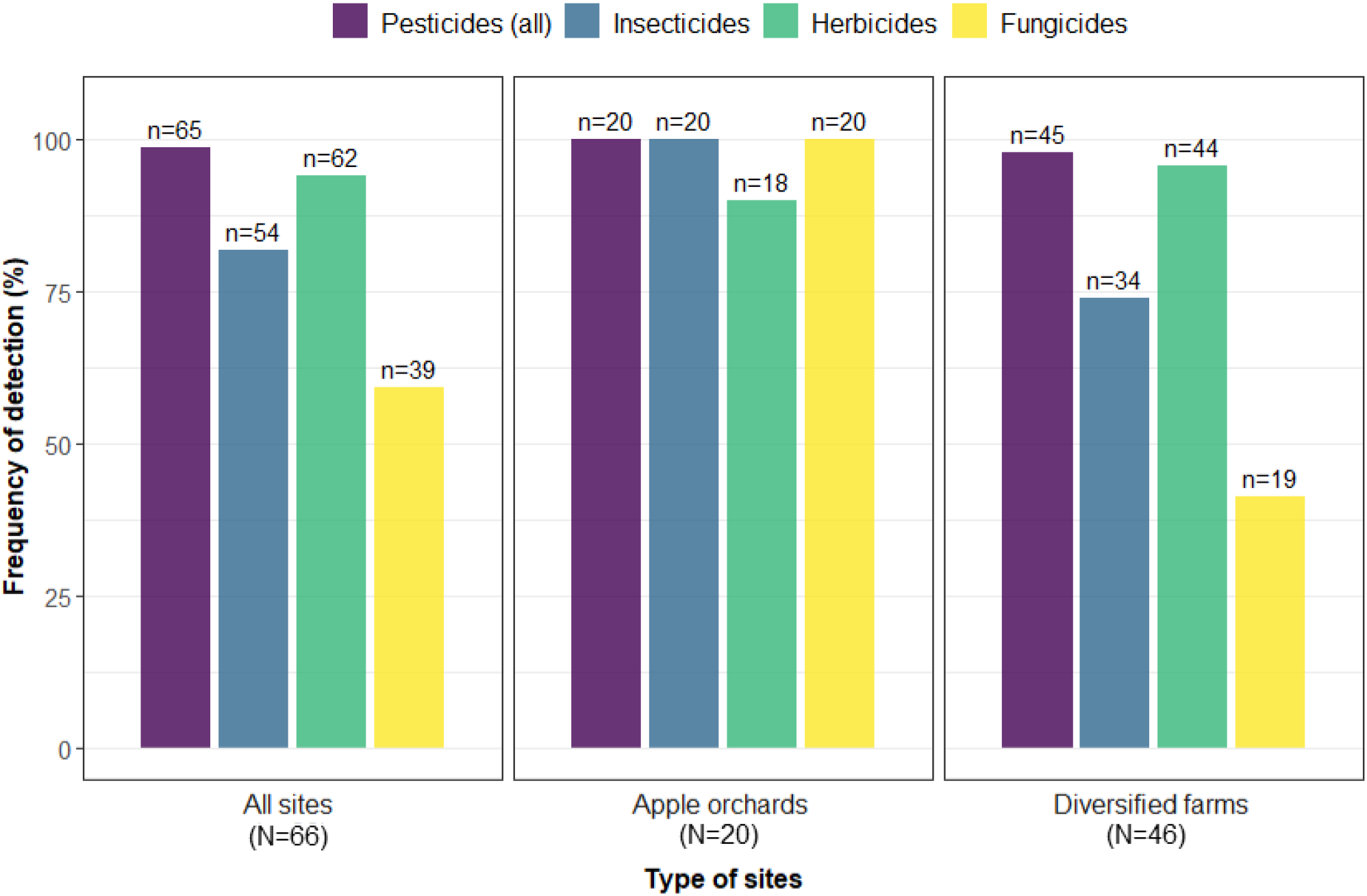
Frequency of soil samples in which at least one pesticide, insecticide, herbicide, or fungicide has been detected. N = number of tested samples; n = number of soils containing pesticide residues.

A total of 53 different AIs were detected in soils, including 27 fungicides, 13 insecticides, and 13 herbicides (Table S3). The pesticide synergist piperonyl butoxide was also found in one sample, along with 11 other AIs. Forty-nine AIs were detected in orchard soils, compared to 29 in soils from diversified farms. The most common AIs detected in samples (Fig. 2) were also retrieved at the highest concentrations (Table S3). Overall, the highest maximum concentration was found for the fungicide boscalid (>1,000 ng/g), while the highest herbicide and insecticide concentrations were found for metolachlor (173.09 ng/g) and chlorantraniliprole (171.31 ng/g), respectively (Table S3). Across both years and cropping systems, the average total concentration of all detected pesticides in soils was 75.9 ng/g ± 260.0 (mean ± SD) and was dominated by fungicide residues (74.8% of pesticide content), followed by insecticide (18.2%) and herbicide (7.0%) residues.

**Figure 2.**
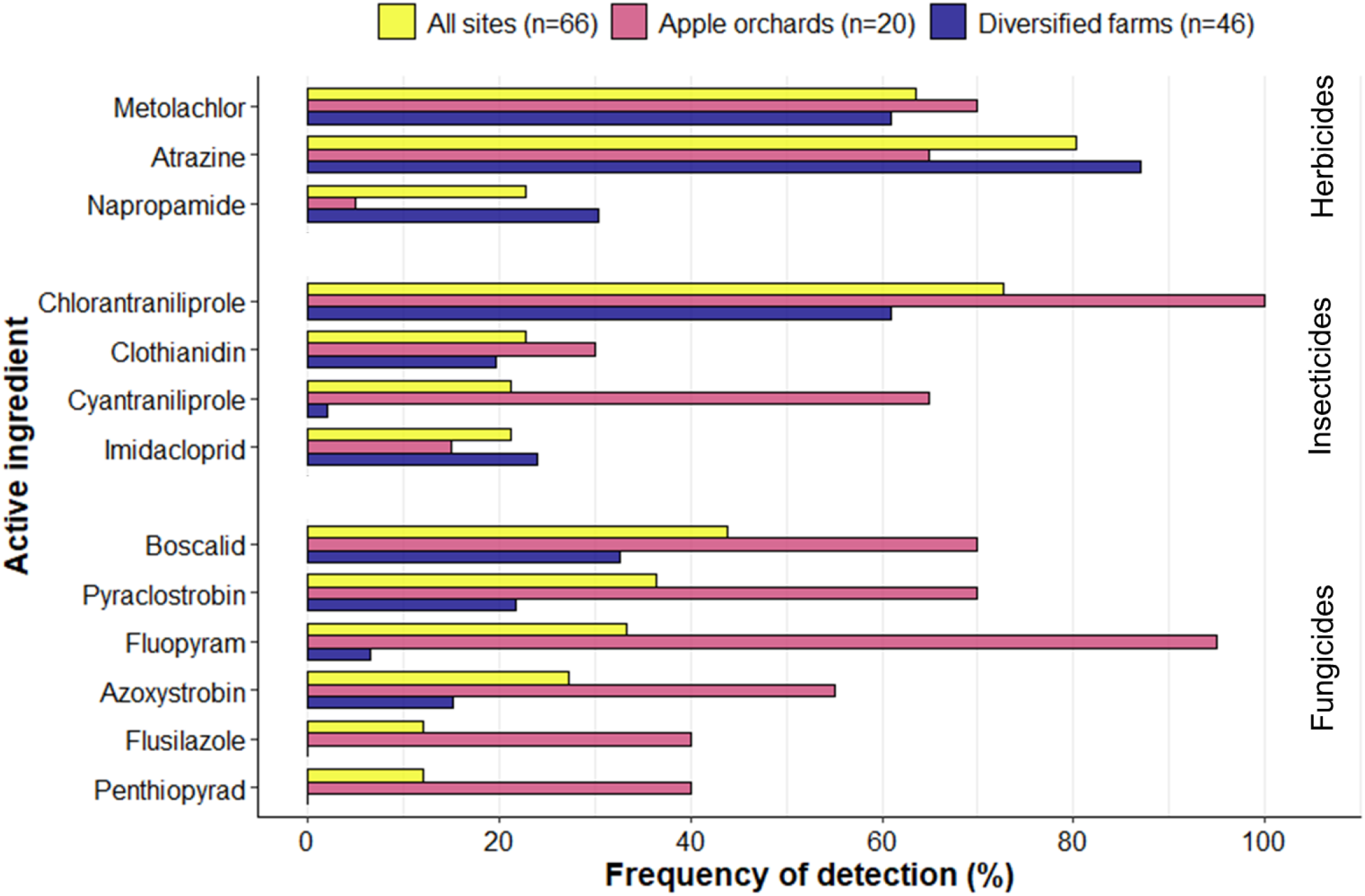
Frequency of detection of the most common AIs detected in soils, per type of cropping system.

Up to 29 different AIs were detected per soil sample, with an average of 12.3 and 4.3 residues in orchard and diversified farm soils, respectively (Fig. 3). Ninety-one percent (60/66) of all soil samples contained residues of at least two AIs and 21.2 % (14/66) contained residues of at least ten AIs. The most common mixtures in soils were atrazine + chlorantraniliprole (57.6% of samples), atrazine + metolachlor (51.5%), and atrazine + chlorantraniliprole + metolachlor (37.9%), often in combination with other pesticide residues (Table S4).

**Figure 3.**
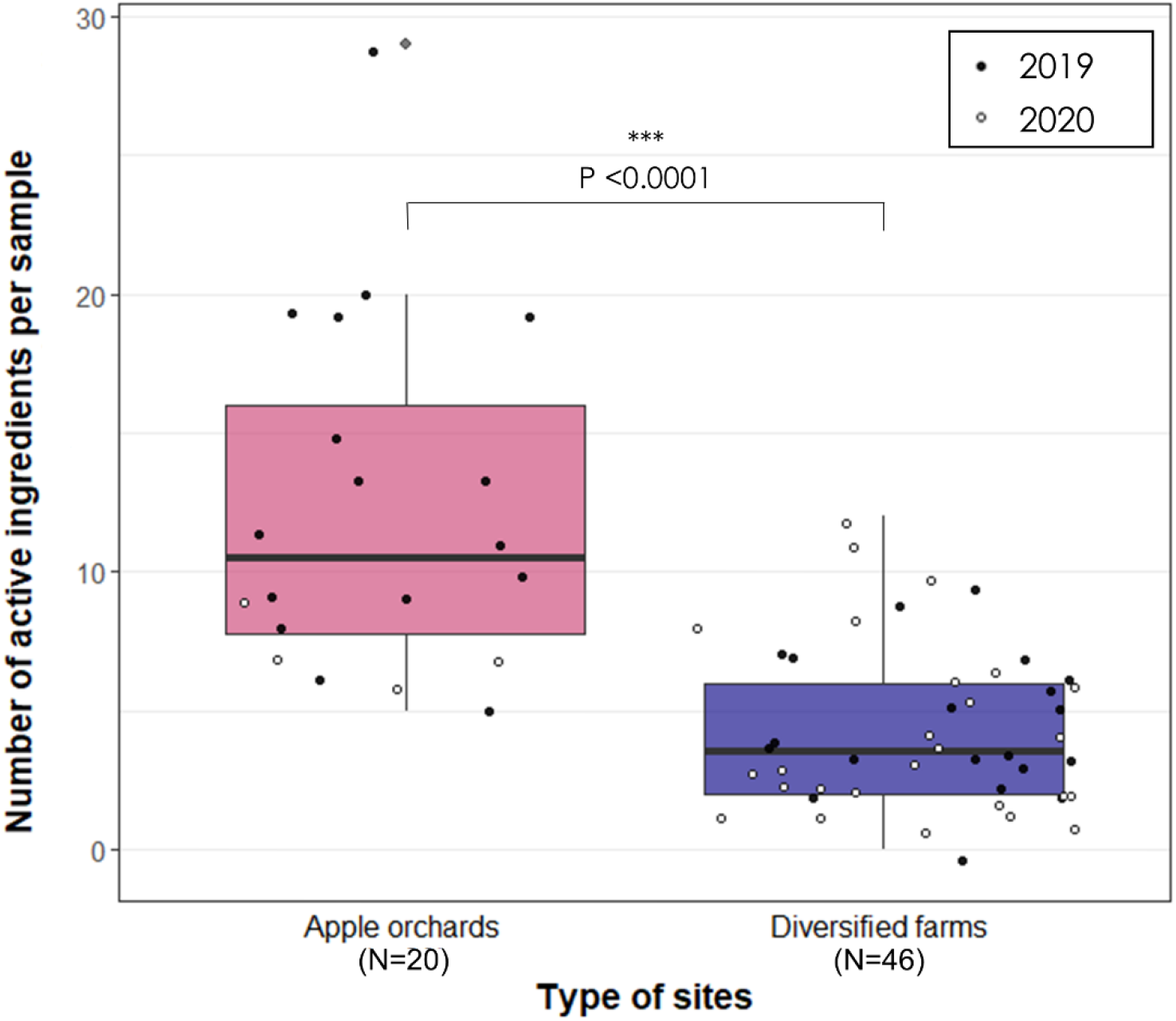
Number of active ingredients detected in agricultural soils sampled at sites identified as possible hibernation sites for bumblebee queens in Ontario, Canada.

### Comparisons between cropping systems

The total pesticide concentration in soils was significantly higher in samples collected from apple orchards compared to those collected from diversified farms (F_(1, 61)_=18.17, P <0.0001) and increased as the distance to the nearest crop decreased (F_(1, 61)_=10.55, P=0.002) in both systems. Orchard soils had a median and maximum total pesticide content of 37.8 and 1,504.2 ng/g respectively, compared to 2.2 and 109.2 ng/g for diversified farm soils (Fig. 4).

**Figure 4.**
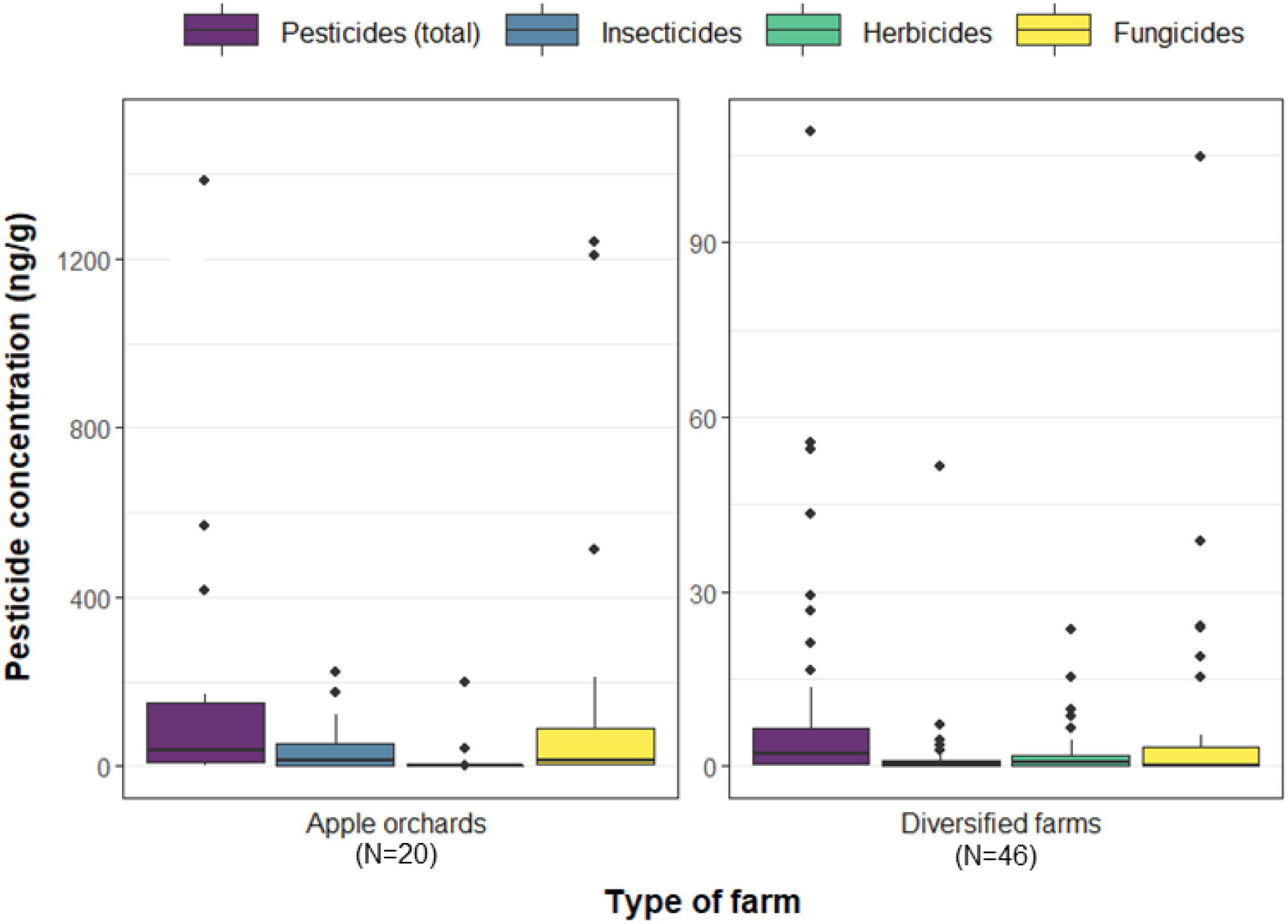
Total residue concentration of pesticides detected in soil samples (i.e., cumulative concentration of all active ingredients), per pesticide group and type of cropping system.

Orchard soils also contained significantly higher numbers of AIs (F_(1, 62)_=26.77, P <0.0001) compared to soils collected from diversified farms. Mixtures of moderate or large numbers (5-29) of residues prevailed in orchard soils relative to mixtures of only a few (0-12) residues for diversified farms (Fig. 3). Besides, 95% (19/20) of soil samples collected in orchards contained at least one insecticide, one herbicide, and one fungicide, compared to 41.3% (19/46) for diversified farms (Table S4).

### Correlations between pesticide residues and soil and pesticide properties

Both soil and pesticide properties influenced the pesticide content in soils (Fig. 5, Table S5), with the persistence of the pesticides in soil (DT50) showing a strong positive correlation with the frequency of detection and average and maximum pesticide concentrations (Fig. 5b). Most (57.8%) of the AIs detected in soils were moderately persistent (DT50: 30-99 days) or persistent (DT50: 100-365 days), while 25% were non persistent (DT50: <30 days) and 17.3% were very persistent (DT50: >365 days). Significant weak negative correlations were also found between pesticide content and both the OM content in soil and slope inclination (Fig. 5a, Table S5). Most soils were sandy loams (57.6%), with a few being loams (19.7%), loamy sands (9.1%), clay loams (9.1%), sandy clays (3.0%), and sands (1.5%).

**Figure 5.**
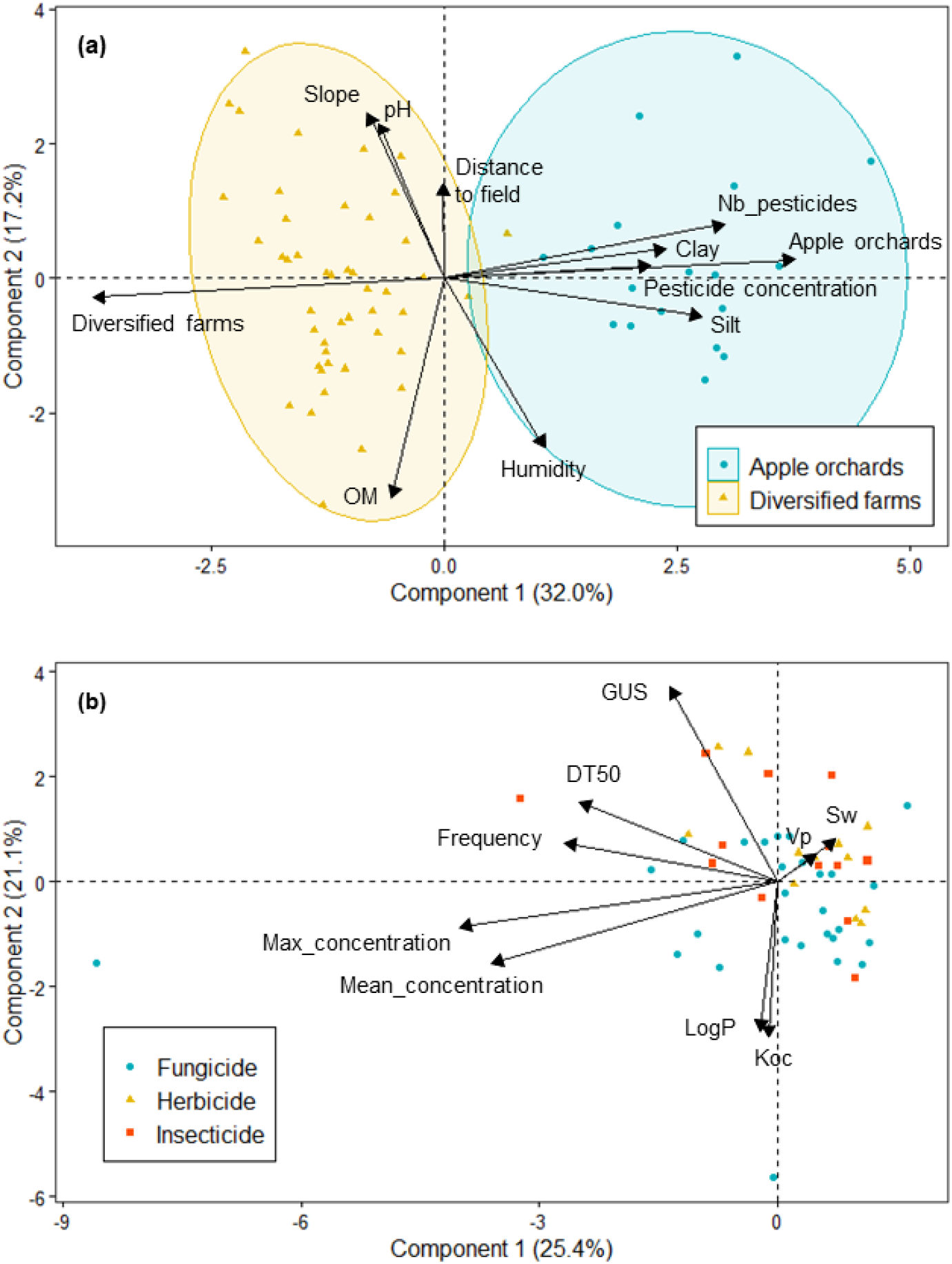
Principal component analysis of pesticide content in soil and soil, site, and pesticide properties. In (a), the total pesticide concentration and number of AIs per sample (n=66) are represented along basic soil properties, slope inclination at sampling site, and distance to the nearest field for each cropping system (apple orchards or diversified farms). In (b), the frequency of detection and the maximum and average concentrations of individual pesticides detected in soil (n=52) are related to their pesticide properties. OM = organic matter content; GUS = leaching potential index; DT50 = soil half-life time; LogP = octanol-water partition coefficient; Koc = organic carbon-water partition coefficient; Sw = solubility in water; Vp = vapor pressure.

### Pesticide residues from confirmed hibernation site

During our 2020 field observations, we were only able to find one bumblebee hibernation site, confirmed by the presence of a bumblebee queen (*B. impatiens*) digging a hibernaculum in the ground. The queen was found at one of the diversified farms, digging in a pile of topsoil (∼4m high) on which scattered weeds had grown. The slope was facing north-west with an inclination of 35°. The soil sampled at this site was a sandy loam (78% sand, 11% silt, 11% clay) and contained low residue levels of four different AIs: the herbicides atrazine (<0.06 ng/g), metolachlor (0.72 ng/g), and napropamide (0.29 ng/g) and the insecticide imidacloprid (<0.3 ng/g). Residue data for this site were included in all data analyses.

## DISCUSSION

Our results show that when hibernating in agricultural areas, bumblebee queens are very likely to be exposed to a wide range of insecticides, fungicides, and herbicides. Pesticide contamination of soils at suitable hibernation sites was found to be widespread, with AIs detected at the highest frequency being broad-spectrum agricultural herbicides (atrazine and metolachlor), insecticides (chlorantraniliprole), and fungicides (boscalid, pyraclostrobin, fluopyram). Overall, fungicides contributed the most to the total pesticide concentrations in soil, while herbicides were the most frequently detected.

From all detections, insecticide residues are expected to be the most concerning for bumblebee health since they are designed to kill insects. The main insecticides detected in soil were the diamides, chlorantraniliprole and cyantraniliprole, and the neonicotinoids, clothianidin and imidacloprid. Neonicotinoid insecticides are well known for their negative effects on bees and other wildlife (Blacquière *et al*. 2012; Goulson 2013), and their use has recently become more restricted in Ontario (OMAFRA 2015) and across Canada (Health Canada 2020). As a result, the use of diamide insecticides has more than doubled since 2015, with more than 100,000 kg of cyantraniliprole and 10,000 kg of chlorantraniliprole sold in Canada in 2019 (Health Canada 2021). While chlorantraniliprole has low acute oral toxicity to both honeybees and bumblebees (Dinter *et al*. 2010; Mundy-Heisz *et al*. 2022), cyantraniliprole is highly toxic to honeybees when exposed by contact (Lewis *et al*. 2016) and possible effects of this insecticide for other bee species are currently unknown. Overall, the insecticide concentrations measured in our samples were low and unlikely to cause acute bumblebee queen mortality, but these concentrations may result in sublethal effects, especially when considering chronic exposure over several months. For instance, the maximum residue concentration found for cyantraniliprole (148 ng/g) represents an exposure to 15.8% of the honeybee contact LD_50_ (934 ng/g) for this AI, close to the 20% threshold set by EFSA (2013) for acute contact exposure level of concern. However, this calculation is imperfect considering that contact transfer factors from soil residues are currently unknown (Thompson 2021). Oral exposure may also be of concern, considering that bumblebee queens use their mandibles to dig into the ground (Alford 1969) and exposure to 148 ng/g of cyantraniliprole represents 13.5% of the honeybee oral LD_50_ (1,100 ng/g), exceeding the 3% EFSA (2013) level of concern for chronic oral exposure. To date, possible impacts of such exposure on bumblebee queens remain unknown.

Compared to insecticides, fungicides and herbicides alone are relatively less toxic to bees, but both groups may interact with other pesticides or ‘inactive’ ingredients included in spray formulations (Siviter *et al*. 2021) or have indirect impacts on bumblebees (Belsky & Joshi 2020). For instance, demethylation inhibitor (DMI) fungicides like those found in our soil samples (difenoconazole, fenbuconazole, flusilazole, metconazole, myclobutanil, propiconazole, triadimenol) can increase the toxicity of pyrethroid and neonicotinoid insecticides (synergism) for bumblebees by inhibiting enzymatic detoxification in insects (Raimets *et al*. 2018). Boscalid, the most commonly detected fungicide in our soils, has also been found to double the acute toxicity of thiamethoxam in honeybees (Tsvetkov *et al*. 2017). This highlights the need for understanding how exposure to these mixtures in soil during underground hibernation may affect bumblebee queens, including whether synergistic effects can occur. The omnipresence and diversity of AI detections and the complexity of pesticide mixtures found in soils at bumblebee hibernation sites (Table S4), and particularly in apple orchards, also deserve more attention.

The diversity of AIs detected in soils is in line with the pesticides approved for use per crop type in Ontario. For instance, many fungicides commonly detected in orchards soils (e.g., penthiopyrad, pyrimethanil) are widely used in apple production, while herbicides such as napropamide are widely applied in vegetable and field crop production (Farm and Food Care Ontario 2015). On the other hand, the most common AIs in soils (e.g., atrazine, boscalid, chlorantraniliprole, metolachlor, pyraclostrobin) were broad-spectrum pesticides that can be applied to a variety of crops. The high occurrence of these pesticides in agricultural soils can also be explained by their moderate-to-high persistence in soil. However, as information on pesticide application at farms involved in this study is not available, we cannot draw clear conclusions between the diversity of products/AIs used on farms and the residues detected in soils.

While previous studies have quantified pesticide content in agricultural field soils, we are the first to consider an exposure specific to hibernating bumblebee queens. This makes comparisons between studies difficult considering that detections are highly dependent on the number and type of residues analysed and that most existing studies only quantified a few AIs in field soils or were confined to one or a few crops. For instance, the analytical methods used in this study did not test for pyrethroid insecticide or glyphosate residues. Considering that glyphosate has been the most widely sold and applied herbicide in Canada in the past decade (Health Canada 2021), its presence in soils at bumblebee hibernation sites is very likely. Residue concentrations in crop soils also largely depend on sampling time, regions, crop type, frequency and intensity of precipitation, agricultural practices, and soil characteristics (Wan *et al*. 2006; Silva *et al*. 2019). For instance, pesticide concentrations in crop field soils have been found to be higher at the start of the growing season compared to later in the summer or fall (Wan *et al*. 2006; Zhou *et al*. 2021). This further emphasizes the importance of considering a sampling timing relevant to the ecology of the species under study, as we did.

The concentrations of atrazine and metolachlor in soils were much lower in the present study than reported for crop soils from Western Canada (Wan *et al*. 2006). Similarly, neonicotinoid concentrations detected in this study were lower than those observed in Ontario corn (Schaafsma *et al*. 2015; Limay-Rios *et al*. 2016; Gaudreault 2020), soybean (Gaudreault 2020), and cucurbit (Willis Chan *et al*. 2019) field soils, but somewhat similar to provincial screening concentrations of agricultural field soils collected during fall (MOECC 2016). Such a trend for lower exposure levels at hibernation sites compared to within fields was expected. We observed a significant tendency of soil pesticide content to decrease as the distance from the nearest crop field increases. Apart from the distance to crop fields, other site characteristics may also provide certain protection against pesticide exposure. For instance, tree cover can reduce residue drift from neighbouring fields (Felsot *et al*. 2011) and potentially decrease pesticide accumulation in hibernation site soil. The slope of the ground may also increase water and pesticide runoff, carrying pesticides off the site, while leaching potential is increased in sandy soils (Pérez-Lucas *et al*. 2019).

Residue concentrations measured in this study likely represent maximum exposure levels for bumblebee queens in Ontario, as soils were collected at the beginning of their hibernation period (Plath 1927) and residue levels are expected to decrease over time (Chaplain *et al*. 2011). However, these residue levels are average concentrations across the top 10 cm soil layer and may, as such, underestimate exposure for bumblebee queens that hibernate at shallower soil depths. This is because pesticide residues often accumulate on the topsoil surface, reaching up to two to three times higher concentrations in the top 1-2 cm of soil surface layer than deeper in the profile (Limay-Rios *et al*. 2016; Silva *et al*. 2019), and bumblebee queens may hibernate as little as 2 cm underground (Hobbs 1967; Alford 1969). Moreover, some queens may choose to hibernate closer to crop fields, leading to potentially increased exposure.

Our results indicate that pesticide exposure in soil varies among cropping systems. In our study, soil samples collected from apple orchards had the highest frequencies of detections, pesticide concentrations, and numbers of pesticides per sample. Similar results have been found in large-scale monitoring studies on soil contamination (Silva *et al*. 2019; Tang & Maggi 2021), in which soils from permanent crops, including orchards, also contained the highest pesticide content and number of detectable pesticides among all tested cropping systems. This is in line with highly intensive pesticide usage in apple orchards and other permanent crops (Mallinger *et al*. 2015; OECD 2019). In Ontario, pesticide use per hectare (and fungicide use more specifically) is the most intensive for fruit production, followed by vegetables and field crops (Farm and Food Care Ontario 2015). Our results suggest that intensive pesticide use can make apple orchards a risky environment for bumblebees and other wild pollinators.

In Canadian apple orchards, landscape enhancements such as hedgerows and flower strips have been linked to an increase in bumblebee abundance and richness (Gervais *et al*. 2021). Such landscape enhancements can provide food resources and nesting and hibernating opportunities for bumblebees, as well as possible shelter from pesticide exposure (Felsot *et al*. 2011; Gervais *et al*. 2021). However, our study shows that these enhancements may not completely protect bumblebees from exposure to pesticide residues in soil if queens hibernate in these areas. Many soil samples analysed in this study were sampled from hedgerows, riparian strips, or edges of forested areas.

The environmental behaviour of pesticides in soil can be related to soil texture (% clay) and OM content, which can both influence pesticide sorption and degradation (Chaplain *et al*. 2011). In our study, we found a significant weak negative correlation between total pesticide content (concentration and number of AIs) and soil OM content. Significant negative correlations with OM content were also found for boscalid and clothianidin (Table S5). The OM content in soil has often been reported to correlate positively with pesticide content (Silva *et al*. 2019; Zhou *et al*. 2021), as most pesticides tend to sorb to OM (Chaplain *et al*. 2011). However, adsorption alone does not dictate pesticide fate in the environment (Chaplain *et al*. 2011). Other factors and processes, such as desorption, degradation, and the physico-chemical characteristics of soil also have a very strong influence on pesticides’ behaviour in soil (Kah *et al*. 2007; Chaplain *et al*. 2011). For instance, the degradation rates of many pesticides, including clothianidin (Li *et al*. 2018), are known to increase with greater OM content in agricultural soils (Kah *et al*. 2007). This is because soils that are high in OM also tend to have a higher diversity and population of soil organisms that can metabolize pesticides, contributing to degradation (Kah *et al*. 2007).

Considering that soils in this study were collected in fall, some pesticide biodegradation by microorganisms likely occurred before sampling, which may explain the lowest content of pesticides in soils with higher OM content.

In addition to confirming that bumblebee queens can indeed hibernate on farms, our observation of a queen digging a hibernaculum in the ground and further quantification of residues at this site support the relevance of our exposure data. Indeed, the pile of topsoil in which we observed a queen digging its underground burrow shared many of the characteristics (slope exposure and inclination, vegetation, soil texture) that we used to identify suitable hibernations sites. Moreover, the identity, number, and concentrations of pesticides detected in the soil sampled at this site approximate the median values measured in diversified farm soils. This strengthens the assertion that bumblebee queens hibernating in agricultural soils are very likely to be exposed to the pesticide AIs and concentrations detected in this study.

## CONCLUSIONS

We generated ecologically relevant exposure estimates of overwintering bumblebee queens to pesticide residues in soil. The occurrences, concentrations, and numbers of AIs detected in soils varied among cropping systems and were the highest in apple orchard soils. Our results support the need for empirical testing of the potential effects of pesticide residues in soils, including co-occurrences, for hibernating bumblebee queens. In future studies, the exposure data we generated can be used to design realistic experiments on the impact of pesticides residues in soil on the health and fitness of bumblebee queens. The differences in exposure that we detected between cropping systems can also be used to better inform regulations that govern the use of agricultural pesticides, notably in apple orchards. Additionally, further exposure studies, conducted across diverse agricultural landscapes and temperate regions, would help develop a better understanding of conditions that lead to significant pesticide exposure for bumblebee queens and how to avoid them.

## AUTHORS CONTRIBUTION

SR and NER conceived the idea and designed the field study; SR collected and analyzed the data and drafted the manuscript; NB and SM performed pesticide residue analysis. All authors contributed to manuscript revisions and approved the final draft for publication.

## ACKNOWLEDGEMENTS

We thank Sébastien Rousseau and Caroline Strang for help with data collection, Susan Willis Chan and Leah Blechschmidt for assistance with site selection, David Sossa for help with sample preparation, and Ryan Prosser for their advice on study design and sharing of lab equipment. This work was supported by an Ontario Ministry of Environment and Climate Change (MOECC) Best in Science grant BIS201617-06, Natural Sciences and Engineering Research Council (NSERC) Discovery Grants 2015-06783 and 2021-04210, and the Food from Thought: Agricultural Systems for a Healthy Planet Initiative, by the Canada First Research Excellence Fund (grant 000054), and the USDA National Institute of Food and Agriculture grant 2018-08603. S.R. is supported by graduate scholarships from The Arrell Food Institute at the University of Guelph and the Fonds de recherche du Québec – Nature et technologies (FRQNT). N.E.R. is supported as the Rebanks Family Chair in Pollinator Conservation by the Weston Family Foundation.

## Notes

### Competing Interest Statement

The authors have declared no competing interest.

