## Supplementary material for "Quantifying exposure of bumblebee (*Bombus* spp.) queens to pesticide residues when hibernating in agricultural soils"

Sabrina Rondeau[a*], Nicolas Baert[b], Scott McArt, Nigel E. Raine[a]

[a] School of Environmental Sciences, University of Guelph, Ontario, Canada
[b] Department of Entomology, Cornell University, Ithaca, New York, USA

**APPENDIX S1.** Detailed methods for the determination of pesticide residues.

**Chemicals and Reagents**

Acetonitrile and water of HPLC grade were purchased from EMD Millipore. LC-MS grade formic acid was purchased from Thermo Scientific. The 5M ammonium formate solution, the QuEChERS extraction packets (4 g MgSO4; 1 g NaCl; 1 g sodium citrate tribasic dihydrate; 0.5 g sodium citrate dibasic sesquihydrate) and the d-SPE kits (150 mg MgSO4 and 25 mg PSA) were purchased from Agilent Technologies.

**Sample preparation**

Frozen soil sample were extracted using the EN 15662 QuEChERS procedure [1] and screened for 247 pesticides (including some metabolites and breakdown products) by liquid chromatography mass spectrometry (LC-MS/MS). Frozen soil samples (10 g) were let to thaw for 1 hour at room temperature then mixed with 10 mL of acetonitrile. After thorough homogenization, 6.5 g of EN 15662 salts were added (4 g MgSO4; 1 g NaCl; 1 g sodium citrate tribasic dihydrate; 0.5 g sodium citrate dibasic sesquihydrate). Samples were vortexed then centrifuged at 7300 × g for 10 minutes. One milliliter of supernatant was collected and transferred into a dispersive solid phase extraction (d-SPE) tube containing 150 mg $MgSO_4$ and 25 mg PSA. After the d-SPE step, 200 µL of supernatant were filtered through 0.22 µm PTFE filter and placed into the autosampler (6°C) for immediate analysis.

**Liquid Chromatography and Mass Spectrometry**

The analysis was performed with a Vanquish Flex UHPLC system (Dionex Softron GmbH, Germering, Germany) coupled with a TSQ Quantis mass spectrometer (Thermo Scientific, San Jose, CA). The UHPLC was fitted with an Acquity UPLC BEH C18 column (100 mm × 2.1 mm, 1.7 µm particle size). The mobile phase consisted of (A) Water containing 2 mM ammonium formate and 0.1% formic acid and (B) Acetonitrile/Water (98:2, v/v) with 2 mM ammonium formate and 0.1% formic acid. The temperature of the column was set at 40°C and the flow rate of the LC was 300 µL/min. The elution program was as follows: 1.5 min equilibration (0% B) prior to injection, 0-0.5 min (0% B, isocratic), 0.5-10 min (0%→70% B, linear gradient), 10-12 min (70%→100% B, linear gradient), 12-15 min (100% B, column wash), 15-15.2 min (100%→0% B, linear gradient), 15.2-17 min (0% B, re-equilibration). The flow from the LC was directed to the mass spectrometer through a Heated Electrospray probe (H-ESI). The settings of the H-ESI were: spray voltage 2500 V for positive mode and 2000 V for negative mode, Sheath gas 55 (arbitrary unit), Auxiliary gas 25 (arbitrary unit), Sweep gas 1 (arbitrary unit), Ion transfer tube temperature 325°C, Vaporizer temperature 350°C.

The MS/MS detection was carried out using the Selected Reaction Monitoring (SRM) mode. Two transitions were monitored for each compound: one for quantification and the other for confirmation. The SRM parameters for each individual compound are summarized in Table S1. The resolution of both Q1 and Q3 was set at 0.7 FWHM, the cycle time was 0.4 s and the pressure of the collision gas (argon) was set at 2 mTorr.

**References:** [1] Foods of plant origin - Multimethod for the determination of pesticide residues using GC- and LC-based analysis following acetonitrile extraction/partitioning and clean-up by dispersive SPE - Modular QuEChERS-method. BS EN 15662:2018.

**Table S1.** List of the 247 pesticides screened for, along with levels of detection (LOD), levels of quantification (LOQ), retention times, and optimized SRM acquisition parameters (RT: Retention time, CE: Collision Energy).

| Compound | LOD (ng/g) | LOQ (ng/g) | RT (min) | RF Lens (V) | Polarity | Precursor (*m/z*) | Quantifying ion (*m/z*) | CE 1 (V) | Confirming ion (*m/z*) | CE 2 (V) |
|---|---|---|---|---|---|---|---|---|---|---|
| Chlormequat chloride | 0.1 | 0.3 | 0.98 | 76 | Positive | 122 | 58 | 27 | 63 | 22 |
| Mepiquat chloride | 0.06 | 0.18 | 1.02 | 83 | Positive | 114 | 98 | 28 | 58 | 26 |
| Cyromazine | 0.5 | 1.5 | 2.65 | 97 | Positive | 167 | 125 | 18 | 85 | 19 |
| Methamidophos | 0.3 | 0.9 | 2.68 | 73 | Positive | 142 | 93.9 | 14 | 124.9 | 14 |
| Acephate | 0.5 | 1.5 | 3.36 | 66 | Positive | 184.1 | 143 | 10 | 94.9 | 23 |
| Pymetrozine | 0.1 | 0.3 | 3.39 | 100 | Positive | 218.1 | 105 | 20 | 78 | 38 |
| Aminocarb | 0.06 | 0.18 | 3.60 | 85 | Positive | 209.1 | 152.1 | 14 | 137 | 24 |
| Formetanate hydrochloride | 0.1 | 0.3 | 3.61 | 114 | Positive | 222.1 | 165.1 | 15 | 92.9 | 35 |
| Omethoate | 0.06 | 0.18 | 3.64 | 81 | Positive | 214 | 182.9 | 12 | 124.9 | 23 |
| Butocarboxim sulfoxide | 0.5 | 0.15 | 3.67 | 77 | Positive | 207 | 132 | 10 | 87.9 | 10 |
| Propamocarb | 0.06 | 0.18 | 3.82 | 89 | Positive | 189.1 | 102 | 17 | 74 | 25 |
| Dinotefuran | 0.1 | 0.3 | 3.87 | 67 | Positive | 203.1 | 113.1 | 10 | 129 | 12 |
| Thiabendazole | 0.05 | 0.15 | 4.30 | 130 | Positive | 202 | 175 | 26 | 131 | 33 |
| Oxamyl | 0.08 | 0.24 | 4.38 | 70 | Positive | 237 | 72.1 | 10 | 90 | 10 |
| Fuberidazole | 0.02 | 0.06 | 4.51 | 126 | Positive | 185.1 | 157 | 22 | 156 | 29 |
| Monocrotophos | 0.08 | 0.24 | 4.52 | 87 | Positive | 224.1 | 193 | 10 | 126.9 | 16 |
| Methomyl | 0.08 | 0.24 | 4.55 | 46 | Positive | 163 | 87.9 | 10 | 106 | 10 |
| Pirimicarb desmethyl | 0.06 | 0.18 | 4.61 | 93 | Positive | 225.1 | 168 | 15 | 72.1 | 21 |
| Butoxycarboxim | 0.02 | 0.06 | 4.75 | 86 | Positive | 223 | 166 | 14 | 105.9 | 10 |
| Dicrotophos | 0.02 | 0.06 | 4.86 | 93 | Positive | 238.1 | 193 | 10 | 112 | 12 |
| Demeton-S-methylsulfone | 0.02 | 0.06 | 4.88 | 110 | Positive | 263 | 168.9 | 16 | 108.9 | 28 |
| Thiamethoxam | 0.05 | 0.15 | 5.02 | 87 | Positive | 292 | 211.1 | 10 | 181 | 22 |
| Ethiofencarb sulfoxide | 0.1 | 0.3 | 5.06 | 75 | Positive | 242.1 | 107 | 18 | 185 | 10 |
| Methiocarb sulfoxide | 0.01 | 0.03 | 5.30 | 89 | Positive | 242.1 | 185 | 14 | 122 | 29 |
| Trichlorfon | 0.5 | 0.15 | 5.31 | 105 | Positive | 256.9 | 108.9 | 18 | 220.9 | 11 |
| Vamidothion | 0.02 | 0.06 | 5.32 | 82 | Positive | 288 | 146.1 | 13 | 118.1 | 23 |
| Schradan | 0.02 | 0.06 | 5.40 | 106 | Positive | 287.1 | 242 | 14 | 135 | 26 |
| Metamitron | 0.1 | 0.3 | 5.41 | 101 | Positive | 203.1 | 175 | 17 | 104 | 24 |
| Clothianidin | 0.2 | 0.6 | 5.48 | 78 | Positive | 250 | 169 | 13 | 131.9 | 17 |
| Ethiofencarb sulfone | 0.1 | 0.3 | 5.52 | 97 | Positive | 258 | 107 | 16 | 200.9 | |
| Fenuron | 0.06 | 0.18 | 5.55 | 84 | Positive | 165 | 72.1 | 16 | 46 | 15 |
| Mevinphos | 0.1 | 0.3 | 5.57 | 82 | Positive | 225 | 193 | 10 | 127 | 17 |
| 3-Hydroxy-carbofuran | 0.1 | 0.3 | 5.58 | 94 | Positive | 238.1 | 181 | 10 | 163 | 16 |
| Pirimicarb | 0.02 | 0.06 | 5.62 | 108 | Positive | 239.1 | 182.1 | 16 | 72 | 22 |
| Chloridazon | 0.1 | 0.3 | 5.64 | 152 | Positive | 222 | 104 | 23 | 77 | 33 |
| Imidacloprid | 0.1 | 0.3 | 5.72 | 94 | Positive | 256 | 209 | 16 | 175 | 18 |

| | | | | | | | | | | |
|---|---|---|---|---|---|---|---|---|---|---|
| Dimethoate | 0.02 | 0.06 | 5.88 | 75 | Positive | 230 | 199 | 10 | 124.9 | 22 |
| Acetamiprid | 0.05 | 0.15 | 6.02 | 92 | Positive | 223 | 125.9 | 22 | 90 | 34 |
| Tricyclazole | 0.02 | 0.06 | 6.09 | 104 | Positive | 190 | 163 | 22 | 136 | 28 |
| Flumetsulam | 0.1 | 0.3 | 6.26 | 136 | Positive | 326 | 129 | 27 | 109 | 51 |
| Methiocarb sulfone | 0.05 | 0.15 | 6.34 | 94 | Positive | 258 | 122 | 19 | 201 | 10 |
| Flupyradifurone | 0.2 | 0.6 | 6.43 | 110 | Positive | 288.9 | 126 | 19 | 89.9 | 38 |
| Imazethapyr | 0.1 | 0.3 | 6.49 | 122 | Positive | 290.1 | 245.1 | 21 | 177 | 28 |
| Thiacloprid | 0.05 | 0.15 | 6.64 | 113 | Positive | 253 | 125.9 | 21 | 90 | 36 |
| Simetryn | 0.02 | 0.06 | 6.65 | 133 | Positive | 214.1 | 124 | 20 | 96 | 24 |
| Desmetryn | 0.02 | 0.06 | 6.66 | 126 | Positive | 214.1 | 172 | 18 | 82 | 30 |
| Tebuthiuron | 0.02 | 0.06 | 6.73 | 105 | Positive | 229 | 172 | 18 | 116 | 27 |
| Sulfoxaflor | 1 | 3 | 6.74 | 115 | Negative | 275.9 | 213 | 17 | 261 | 13 |
| Prometon | 0.02 | 0.06 | 6.80 | 118 | Positive | 226.2 | 184.1 | 19 | 142.1 | 23 |
| Carbetamide | 0.08 | 0.24 | 6.83 | 73 | Positive | 237.1 | 192 | 10 | 118 | 16 |
| Hexazinone | 0.02 | 0.06 | 6.84 | 100 | Positive | 253.2 | 171 | 16 | 71.1 | 31 |
| Terbumeton | 0.06 | 0.18 | 6.91 | 118 | Positive | 226.2 | 170 | 17 | 142 | 23 |
| Bromacil | 0.5 | 1.5 | 7.00 | 78 | Positive | 261 | 204.9 | 15 | 187.9 | 28 |
| Thidiazuron | 0.2 | 0.6 | 7.07 | 90 | Positive | 221 | 101.9 | 16 | 94 | 14 |
| Ancymidol | 0.2 | 0.6 | 7.09 | 130 | Positive | 257.1 | 135 | 25 | 81 | 25 |
| Imazaquin | 0.1 | 0.3 | 7.17 | 124 | Positive | 312.1 | 267 | 22 | 199 | 28 |
| Oxadixyl | 0.1 | 0.3 | 7.18 | 62 | Positive | 296.1 | 279 | 10 | 219.1 | 16 |
| Metolcarb | 0.3 | 0.9 | 7.19 | 73 | Positive | 166 | 109 | 10 | 93.9 | 31 |
| Cyanazine | 0.1 | 0.3 | 7.27 | 124 | Positive | 241.1 | 214 | 17 | 104 | 29 |
| Azamethiphos | 0.02 | 0.06 | 7.38 | 108 | Positive | 324.9 | 183 | 16 | 112 | 34 |
| Lenacil | 0.1 | 0.3 | 7.48 | 86 | Positive | 235.1 | 153 | 16 | 136 | 32 |
| Thiophanate-methyl | 0.05 | 0.15 | 7.51 | 119 | Positive | 343 | 151 | 20 | 311 | 10 |
| Methoprotryne | 0.02 | 0.06 | 7.68 | 139 | Positive | 272.1 | 198 | 23 | 170 | 29 |
| Ametryn | 0.02 | 0.06 | 7.69 | 128 | Positive | 228.1 | 186 | 18 | 96 | 26 |
| Malaoxon | 0.02 | 0.06 | 7.70 | 101 | Positive | 315 | 98.9 | 23 | 127 | 12 |
| Imazalil | 0.08 | 0.24 | 7.71 | 135 | Positive | 297 | 158.9 | 24 | 201 | 18 |
| Florasulam | 0.5 | 1.5 | 7.74 | 147 | Positive | 360 | 129 | 24 | 192 | 16 |
| Pyroxsulam | 0.4 | 1.2 | 7.75 | 152 | Positive | 435 | 195 | 27 | 257.9 | 23 |
| Bentazone | 0.1 | 0.3 | 7.76 | 120 | Negative | 238.9 | 132 | 27 | 197 | 21 |
| Fenthion sulfoxide | 0.1 | 0.3 | 7.79 | 117 | Positive | 295 | 279.9 | 19 | 108.9 | 32 |
| Carbofuran | 0.02 | 0.06 | 7.80 | 79 | Positive | 222.1 | 165 | 12 | 123 | 22 |
| Bendiocarb | 0.06 | 0.18 | 7.81 | 81 | Positive | 224.1 | 167 | 10 | 109 | 18 |
| Methabenzthiazuron | 0.02 | 0.06 | 7.84 | 85 | Positive | 222.1 | 165 | 17 | 150 | 33 |
| Chlorotoluron | 0.08 | 0.24 | 7.98 | 95 | Positive | 213.1 | 72 | 18 | 46 | 16 |
| 4-Hydroxy-chlorothalonil | 1 | 3 | 8.03 | 141 | Negative | 244.8 | 181.9 | 29 | 174.9 | 27 |
| Cycluron | 0.08 | 0.24 | 8.04 | 99 | Positive | 199.1 | 89 | 15 | 72.1 | 22 |
| Carbaryl | 0.05 | 0.15 | 8.08 | 73 | Positive | 202 | 145 | 10 | 127 | 29 |

| | | | | | | | | | | |
|---|---|---|---|---|---|---|---|---|---|---|
| Fluometuron | 0.2 | 0.6 | 8.09 | 116 | Positive | 233 | 72 | 19 | 46 | 18 |
| Forchlorfenuron | 0.08 | 0.24 | 8.11 | 106 | Positive | 248 | 129 | 18 | 93 | 33 |
| Sulfentrazone | 2 | 6 | 8.13 | 192 | Negative | 384.9 | 307 | 23 | 199 | 36 |
| Atrazine | 0.02 | 0.06 | 8.14 | 129 | Positive | 216 | 174 | 18 | 104 | 29 |
| Flutriafol | 0.1 | 0.3 | 8.15 | 117 | Positive | 302 | 70 | 19 | 123 | 28 |
| Fosthiazate | 0.02 | 0.06 | 8.16 | 91 | Positive | 284 | 228 | 10 | 103.9 | 21 |
| Bromoxynil | 1 | 3 | 8.21 | 152 | Negative | 275.7 | 78.8 | 30 | 80.8 | 30 |
| Metosulam | 0.2 | 0.6 | 8.26 | 192 | Positive | 417.9 | 174.9 | 26 | 140 | 50 |
| Pyrimethanil | 0.05 | 0.15 | 8.27 | 150 | Positive | 200.1 | 107 | 24 | 82 | 26 |
| Ethiofencarb | 0.5 | 1.5 | 8.30 | 86 | Positive | 226.1 | 107 | 16 | 169 | 10 |
| Monolinuron | 0.3 | 0.9 | 8.31 | 78 | Positive | 215 | 125.9 | 18 | 148 | 15 |
| Ofurace | 0.06 | 0.18 | 8.32 | 115 | Positive | 282.1 | 254.1 | 12 | 160.1 | 24 |
| Isoproturon | 0.06 | 0.18 | 8.34 | 106 | Positive | 207.1 | 72 | 19 | 165 | 14 |
| Metalaxyl | 0.05 | 0.15 | 8.35 | 104 | Positive | 280.1 | 220.1 | 14 | 248.1 | 10 |
| Azaconazole | 0.06 | 0.18 | 8.36 | 133 | Positive | 300 | 158.9 | 28 | 230.9 | 17 |
| Diuron | 0.2 | 0.6 | 8.37 | 95 | Positive | 233 | 72 | 18 | 46 | 17 |
| Dodemorph | 0.02 | 0.06 | 8.38 | 136 | Positive | 282.2 | 116.1 | 21 | 98 | 27 |
| Cyantraniliprole | 0.1 | 0.3 | 8.39 | 137 | Positive | 474.9 | 285.8 | 14 | 443.9 | 19 |
| Griseofulvin | 0.1 | 0.3 | 8.46 | 131 | Positive | 353 | 165 | 20 | 285 | 19 |
| Fensulfothion | 0.06 | 0.18 | 8.60 | 117 | Positive | 309 | 280.9 | 15 | 252.9 | 18 |
| Isoprocarb | 0.1 | 0.3 | 8.61 | 86 | Positive | 194.1 | 94.9 | 15 | 137 | 10 |
| Metobromuron | 0.1 | 0.3 | 8.62 | 99 | Positive | 258.9 | 169.9 | 19 | 148 | 16 |
| Heptenophos | 0.1 | 0.3 | 8.63 | 97 | Positive | 251 | 127 | 14 | 125 | 13 |
| Prometryn | 0.02 | 0.06 | 8.75 | 123 | Positive | 242.1 | 158 | 23 | 200 | 18 |
| Metazachlor | 0.02 | 0.06 | 8.79 | 84 | Positive | 278.1 | 134.1 | 22 | 210 | 10 |
| Spiroxamine | 0..02 | 0.06 | 8.80 | 126 | Positive | 298.2 | 144.1 | 20 | 100 | 30 |
| Terbutryn | 0.02 | 0.06 | 8.84 | 123 | Positive | 242.1 | 186 | 18 | 68 | 40 |
| Ioxynil | 1 | 3 | 8.92 | 127 | Negative | 369.7 | 126.8 | 35 | 214.8 | 32 |
| Dimefuron | 0.06 | 0.18 | 8.97 | 159 | Positive | 339 | 166.9 | 22 | 72.1 | 26 |
| Clomazone | 0.08 | 0.24 | 9.00 | 89 | Positive | 240.1 | 124.9 | 21 | 89 | 47 |
| Chlorantraniliprole | 0.1 | 0.3 | 9.03 | 137 | Positive | 483.9 | 452.9 | 17 | 285.8 | 11 |
| Benodanil | 0.08 | 0.24 | 9.11 | 139 | Positive | 323.9 | 230.9 | 24 | 202.9 | 38 |
| Propazine | 0.02 | 0.06 | 9.12 | 130 | Positive | 230.1 | 188.1 | 18 | 146.1 | 23 |
| Paclobutrazol | 0.1 | 0.3 | 9.20 | 122 | Positive | 294.1 | 70 | 21 | 129.9 | 37 |
| Isocarbophos | 0.3 | 0.9 | 9.26 | 53 | Positive | 307 | 231 | 16 | 273 | 10 |
| Dimethomorph | 0.06 | 0.18 | 9.27 | 164 | Positive | 388.1 | 301 | 20 | 165 | 31 |
| Desmedipham | 0.1 | 0.3 | 9.33 | 118 | Positive | 301 | 182 | 10 | 136 | 21 |
| Phenmedipham | 0.1 | 0.3 | 9.34 | 108 | Positive | 301 | 168 | 10 | 136 | 20 |
| Terbuthylazine | 0.06 | 0.18 | 9.35 | 102 | Positive | 230.1 | 174 | 17 | 96 | 28 |
| Fenobucarb | 0.1 | 0.3 | 9.36 | 86 | Positive | 208.1 | 94.9 | 15 | 151.9 | 10 |
| Methiocarb | 0.2 | 0.6 | 9.37 | 84 | Positive | 226.1 | 169 | 10 | 121 | 19 |

| | | | | | | | | | | |
|---|---|---|---|---|---|---|---|---|---|---|
| Fluazifop | 0.3 | 0.9 | 9.38 | 144 | Positive | 328 | 282 | 19 | 254 | 27 |
| Ethiprole | 1.5 | 4.5 | 9.42 | 147 | Positive | 397 | 350.9 | 21 | 254.9 | 35 |
| Azinphos-methyl | 1.5 | 4.5 | 9.43 | 77 | Positive | 318 | 132 | 15 | 77.1 | 34 |
| Flumioxazin | 5 | 15 | 9.46 | 191 | Positive | 355 | 327 | 21 | 299 | 29 |
| Crotoxyphos | 0.02 | 0.06 | 9.47 | 76 | Positive | 332 | 211 | 10 | 127 | 26 |
| Diethofencarb | 0.1 | 0.3 | 9.49 | 87 | Positive | 268.1 | 226.1 | 10 | 124 | 32 |
| Fludioxonil | 0.3 | 0.9 | 9.53 | 130 | Negative | 246.9 | 180 | 30 | 126 | 32 |
| Dimethametryn | 0.06 | 0.18 | 9.54 | 135 | Positive | 256.1 | 185.9 | 21 | 90.9 | 29 |
| Spirotetramat | 0.06 | 0.18 | 9.55 | 140 | Positive | 374.1 | 302.1 | 17 | 330.2 | 16 |
| Dimethenamid | 0.06 | 0.18 | 9.61 | 89 | Positive | 276.1 | 244 | 13 | 168 | 24 |
| Bromuconazole | 0.3 | 0.9 | 9.62 | 151 | Positive | 377.9 | 159 | 29 | 160.9 | 30 |
| Chloroxuron | 0.1 | 0.3 | 9.63 | 126 | Positive | 291 | 72 | 20 | 46.1 | 17 |
| Phosmet | 1 | 3 | 9.64 | 95 | Positive | 318 | 159.9 | 13 | 133 | 36 |
| Benoxacor | 2 | 6 | 9.65 | 118 | Positive | 260 | 149 | 18 | 134 | 29 |
| Iprovalicarb | 0.04 | 0.12 | 9.66 | 95 | Positive | 321.1 | 119.1 | 20 | 203 | 10 |
| Promecarb | 0.1 | 0.3 | 9.67 | 81 | Positive | 208.1 | 151.1 | 10 | 109 | 17 |
| Chlorbromuron | 1 | 3 | 9.68 | 124 | Positive | 293 | 182 | 16 | 203.9 | 19 |
| Cumyluron | 0.06 | 0.18 | 9.72 | 94 | Positive | 303.1 | 185 | 13 | 124.9 | 34 |
| Simeconazole | 0.2 | 0.6 | 9.73 | 116 | Positive | 294.1 | 70 | 20 | 135 | 21 |
| Azoxystrobin | 0.02 | 0.06 | 9.74 | 127 | Positive | 404 | 372 | 15 | 329 | 31 |
| Fenamiphos | 0.02 | 0.06 | 9.76 | 117 | Positive | 304.1 | 216.9 | 23 | 202 | 35 |
| Cyprodynil | 0.02 | 0.06 | 9.80 | 136 | Positive | 226.1 | 93 | 35 | 108.1 | 27 |
| Myclobutanil | 0.05 | 0.15 | 9.81 | 134 | Positive | 289 | 70 | 20 | 125 | 34 |
| Fenamidone | 0.05 | 0.15 | 9.82 | 112 | Positive | 312 | 236.1 | 14 | 92 | 24 |
| Epoxiconazole | 0.06 | 0.18 | 9.83 | 121 | Positive | 330 | 121 | 22 | 101 | 44 |
| Phoxim | 0.5 | 0.15 | 9.84 | 73 | Positive | 299 | 77 | 29 | 129 | 10 |
| Etaconazole | 0.1 | 0.3 | 9.85 | 136 | Positive | 327.9 | 159 | 29 | 55.1 | 21 |
| Fluxapyroxad | 0.2 | 0.6 | 9.88 | 133 | Positive | 382 | 362 | 15 | 342 | 21 |
| Triadimefon | 0.2 | 0.6 | 9.89 | 111 | Positive | 294 | 197 | 16 | 69 | 21 |
| Mandipropamid | 0.04 | 0.12 | 9.90 | 126 | Positive | 412.1 | 328 | 15 | 356 | 10 |
| Boscalid | 1 | 3 | 9.94 | 150 | Positive | 343 | 307 | 20 | 271 | 32 |
| Propyzamide | 0.5 | 1.5 | 9.96 | 88 | Positive | 256 | 190 | 14 | 172.8 | 22 |
| Fluquinoconazole | 1 | 3 | 9.98 | 139 | Positive | 375.9 | 349 | 19 | 307.1 | 26 |
| Spinosad | 0.06 | 0.18 | 9.99 | 168 | Positive | 732.4 | 142.1 | 29 | 98 | 43 |
| Fenhexamid | 2 | 6 | 10.00 | 136 | Positive | 302 | 97 | 24 | 55 | 34 |
| Bupirimate | 0.06 | 0.18 | 10.01 | 125 | Positive | 317.1 | 166 | 25 | 210.1 | 24 |
| Tetraconazole | 0.2 | 0.6 | 10.06 | 152 | Positive | 372 | 159 | 31 | 70 | 23 |
| Napropamide | 0.02 | 0.06 | 10.09 | 104 | Positive | 272.1 | 199 | 13 | 171 | 19 |
| Fluopicolide | 0.1 | 0.3 | 10.10 | 131 | Positive | 382.9 | 172.8 | 24 | 144.9 | 48 |
| Tebuconazole | 0.2 | 0.6 | 10.13 | 130 | Positive | 308.1 | 70 | 23 | 124.9 | 37 |
| Prochloraz | 0.1 | 0.3 | 10.14 | 99 | Positive | 376 | 307.9 | 11 | 70 | 25 |

| | | | | | | | | | | |
|---|---|---|---|---|---|---|---|---|---|---|
| Isoxaben | 0.06 | 0.18 | 10.15 | 125 | Positive | 333.1 | 165 | 20 | 106.9 | 55 |
| Fluopyram | 0.02 | 0.06 | 10.16 | 126 | Positive | 397 | 208 | 22 | 173 | 29 |
| Trietazine | 0.06 | 0.18 | 10.18 | 129 | Positive | 230.1 | 202 | 17 | 99 | 23 |
| Methoxyfenozide | 0.05 | 0.15 | 10.19 | 77 | Positive | 369.1 | 149 | 17 | 313.1 | 10 |
| Bifenazate | 0.04 | 0.12 | 10.20 | 84 | Positive | 301.1 | 198 | 10 | 170 | 18 |
| Flusilazole | 0.06 | 0.18 | 10.21 | 146 | Positive | 316.1 | 247 | 18 | 165 | 27 |
| Mepronil | 0.04 | 0.12 | 10.25 | 94 | Positive | 270.1 | 119 | 23 | 228.1 | 14 |
| Diflubenzuron | 2 | 6 | 10.26 | 134 | Positive | 311 | 158 | 13 | 141 | 32 |
| Fenbuconazole | 0.1 | 0.3 | 10.29 | 144 | Positive | 337 | 125 | 31 | 70 | 21 |
| Haloxyfop | 5 | 15 | 10.33 | 128 | Positive | 362 | 316 | 18 | 91 | 30 |
| Isoprothiolane | 0.03 | 0.09 | 10.36 | 87 | Positive | 291 | 231 | 10 | 188.9 | 22 |
| Penconazole | 0.1 | 0.3 | 10.38 | 102 | Positive | 284 | 158.9 | 31 | 70 | 18 |
| Bitertanol | 0.5 | 1.5 | 10.39 | 85 | Positive | 338.1 | 269.1 | 10 | 70 | 10 |
| Triazophos | 0.04 | 0.12 | 10.40 | 121 | Positive | 314 | 162.1 | 19 | 119 | 34 |
| Hexaconazole | 0.3 | 0.9 | 10.41 | 129 | Positive | 314 | 70 | 21 | 159 | 32 |
| Metolachlor | 0.1 | 0.3 | 10.45 | 102 | Positive | 284.1 | 252 | 15 | 176.1 | 26 |
| Fenoxycarb | 0.5 | 1.5 | 10.46 | 97 | Positive | 302.1 | 116 | 10 | 88 | 19 |
| Metconazole | 0.2 | 0.6 | 10.47 | 131 | Positive | 320.1 | 70 | 24 | 125 | 39 |
| Alachlor | 0.2 | 0.6 | 10.49 | 82 | Positive | 270.1 | 238 | 10 | 162.1 | 21 |
| Fumagillin | 1 | 3 | 10.50 | 124 | Positive | 459.1 | 131 | 26 | 177 | 14 |
| Flufenacet | 0.2 | 0.6 | 10.53 | 95 | Positive | 364 | 194.1 | 10 | 152 | 19 |
| Fluoxastrobin | 0.02 | 0.06 | 10.54 | 172 | Positive | 459 | 427 | 18 | 188 | 35 |
| Dimoxystrobin | 0.02 | 0.06 | 10.56 | 89 | Positive | 327.1 | 205.1 | 10 | 116 | 22 |
| Azinphos-ethyl | 0.5 | 1.5 | 10.56 | 84 | Positive | 346 | 132 | 1 | 77 | 37 |
| Propetamphos | 2 | 6 | 10.57 | 79 | Positive | 282 | 138 | 17 | 156 | 10 |
| Diniconazole | 0.5 | 1.5 | 10.62 | 149 | Positive | 326 | 70 | 26 | 159 | 32 |
| Propiconazole | 0.2 | 0.6 | 10.63 | 150 | Positive | 342 | 158.9 | 30 | 69.1 | 20 |
| Neburon | 0.1 | 0.3 | 10.64 | 134 | Positive | 275 | 88.1 | 16 | 57.1 | 21 |
| Butafenacil | 0.06 | 0.18 | 10.73 | 133 | Positive | 492 | 330.9 | 23 | 179.9 | 43 |
| Tebufenozide | 0.02 | 0.06 | 10.75 | 82 | Positive | 353.1 | 297.1 | 10 | 133 | 19 |
| Spinetoram | 0.05 | 0.15 | 10.76 | 217 | Positive | 748.3 | 142.1 | 30 | 98 | 44 |
| Triadimenol | 0.02 | 0.06 | 10.77 | 118 | Positive | 297 | 133 | 13 | 105 | 32 |
| Chlorfenvinphos | 0.1 | 0.3 | 10.83 | 121 | Positive | 359 | 155 | 13 | 169.9 | 40 |
| Flubendiamide | 0.5 | 1.5 | 10.87 | 145 | Positive | 682.9 | 407.9 | 10 | 273.9 | 31 |
| Carfentrazone-ethyl | 0.2 | 0.6 | 10.91 | 173 | Positive | 412 | 345.9 | 23 | 365.9 | 17 |
| Cyazofamid | 0.5 | 1.5 | 10.92 | 92 | Positive | 325 | 107.9 | 14 | 261 | 10 |
| Fipronil | 0.1 | 0.3 | 10.96 | 149 | Negative | 434.8 | 330 | 16 | 249.9 | 27 |
| Kresoxim-methyl | 0.1 | 0.3 | 10.97 | 89 | Positive | 314.1 | 267 | 10 | 222.1 | 13 |
| Picoxystrobin | 0.02 | 0.06 | 11.04 | 83 | Positive | 368 | 205 | 10 | 145 | 21 |
| Isoxadifen-ethyl | 0.3 | 0.9 | 11.05 | 112 | Positive | 296.1 | 232 | 17 | 263 | 10 |
| Triflumizole | 0.05 | 0.15 | 11.06 | 90 | Positive | 346 | 278.1 | 10 | 73.1 | 16 |

| | | | | | | | | | | |
|---|---|---|---|---|---|---|---|---|---|---|
| Difenoconazole | 0.05 | 0.15 | 11.07 | 165 | Positive | 406 | 251 | 26 | 337 | 18 |
| Carpropamid | 0.3 | 0.9 | 11.08 | 98 | Positive | 334 | 139 | 19 | 196 | 12 |
| Penthiopyrad | 0.02 | 0.06 | 11.10 | 126 | Positive | 360.1 | 276 | 15 | 256 | 21 |
| Anilofos | 0.02 | 0.06 | 11.21 | 120 | Positive | 367.9 | 198.9 | 14 | 124.9 | 31 |
| Phenthoate | 1 | 3 | 11.25 | 98 | Positive | 321 | 247 | 10 | 274.9 | 10 |
| Etrimfos | 0.02 | 0.06 | 11.26 | 113 | Positive | 293 | 265 | 17 | 124.9 | 26 |
| Zoxamide | 0.1 | 0.3 | 11.29 | 116 | Positive | 336 | 186.9 | 22 | 159 | 39 |
| Coumaphos | 1 | 3 | 11.37 | 127 | Positive | 363 | 227 | 26 | 306.9 | 18 |
| Pyraclostrobin | 0.02 | 0.06 | 11.37 | 114 | Positive | 388.1 | 194 | 13 | 163 | 24 |
| Benzoylprop-ethyl | 0.1 | 0.3 | 11.40 | 102 | Positive | 366 | 105 | 17 | 77 | 46 |
| Fipronil sulfone | 0.06 | 0.18 | 11.50 | 152 | Negative | 450.8 | 414.9 | 17 | 282 | 28 |
| Hexaflumuron | 2 | 6 | 11.51 | 118 | Negative | 458.7 | 439 | 10 | 275.9 | 20 |
| Thiobencarb | 0.3 | 0.9 | 11.52 | 88 | Positive | 258 | 125 | 20 | 89 | 48 |
| Pencycuron | 0.02 | 0.06 | 11.54 | 146 | Positive | 329.1 | 125 | 27 | 218 | 16 |
| Sulfotep | 0.1 | 0.3 | 11.56 | 103 | Positive | 323 | 170.9 | 15 | 96.8 | 36 |
| Pirimiphos-methyl | 0.02 | 0.06 | 11.65 | 112 | Positive | 306 | 164 | 23 | 108 | 30 |
| Cycloate | 0.1 | 0.3 | 11.70 | 79 | Positive | 216 | 154.1 | 13 | 83.1 | 16 |
| Haloxyfop-methyl | 0.1 | 0.3 | 11.71 | 134 | Positive | 376 | 316 | 16 | 91 | 32 |
| Piperophos | 0.02 | 0.06 | 11.75 | 122 | Positive | 354 | 170.8 | 22 | 255 | 14 |
| Benzoximate | 0.02 | 0.06 | 11.76 | 77 | Positive | 364 | 199 | 10 | 105 | 25 |
| Cyflufenamid | 0.1 | 0.3 | 11.77 | 128 | Positive | 413.1 | 295 | 15 | 241 | 23 |
| Indoxacarb | 0.3 | 0.9 | 11.78 | 164 | Positive | 527.9 | 203 | 39 | 150 | 24 |
| Metrafenone | 0.06 | 0.18 | 11.83 | 124 | Positive | 409 | 209 | 15 | 226.9 | 20 |
| Trifloxystrobin | 0.02 | 0.06 | 11.84 | 128 | Positive | 409 | 186 | 18 | 145 | 44 |
| Imibenconazole | 0.3 | 0.9 | 11.85 | 157 | Positive | 410.9 | 125 | 31 | 170.9 | 20 |
| Profenophos | 0.4 | 1.2 | 11.87 | 130 | Positive | 374.9 | 304.8 | 19 | 346.8 | 13 |
| Quizalofop-ethyl | 0.08 | 0.24 | 11.88 | 147 | Positive | 373 | 299 | 19 | 271 | 25 |
| Tebufenpyrad | 0.1 | 0.3 | 11.93 | 137 | Positive | 334.1 | 117 | 36 | 145 | 27 |
| Prosulfocarb | 0.06 | 0.18 | 11.98 | 94 | Positive | 252.1 | 91 | 23 | 128 | 13 |
| Quinoxyfen | 0.06 | 0.18 | 12.04 | 168 | Positive | 308 | 197 | 33 | 162 | 46 |
| Piperonyl butoxide | 0.02 | 0.06 | 12.11 | 92 | Positive | 356.2 | 177 | 12 | 119 | 34 |
| Buprofezin | 0.02 | 0.06 | 12.12 | 99 | Positive | 306.1 | 201.1 | 12 | 116 | 16 |
| Fluazinam | 0.1 | 0.3 | 12.17 | 138 | Negative | 462.8 | 415.9 | 20 | 397.9 | 16 |
| Tetramethrin | 0.3 | 0.9 | 12.18 | 106 | Positive | 332.1 | 164 | 24 | 135.1 | 18 |
| Furathiocarb | 0.02 | 0.06 | 12.19 | 126 | Positive | 383.1 | 195 | 19 | 252 | 13 |
| Sethoxydim | 0.06 | 0.18 | 12.24 | 115 | Positive | 328.2 | 178 | 19 | 282 | 12 |
| Esprocarb | 0.06 | 0.18 | 12.32 | 110 | Positive | 266.1 | 91 | 25 | 71 | 15 |
| Flufenoxuron | 8 | 24 | 12.37 | 126 | Positive | 489 | 158 | 19 | 140.9 | 42 |
| Tralkoxydim | 2 | 6 | 12.42 | 104 | Positive | 330.2 | 284.2 | 13 | 138.1 | 20 |
| Chlorpyrifos | 3 | 9 | 12.44 | 106 | Positive | 349.9 | 197.9 | 20 | 96.9 | 31 |
| Hexythiazox | 0.5 | 1.5 | 12.46 | 107 | Positive | 353 | 228 | 15 | 168 | 25 |

| | | | | | | | | | | |
|---|---|---|---|---|---|---|---|---|---|---|
| Fenpyroximate | 0.05 | 0.15 | 12.51 | 146 | Positive | 422.1 | 366 | 16 | 214.1 | 30 |
| Chlorfluazuron | 8 | 24 | 12.57 | 172 | Positive | 539.9 | 382.9 | 21 | 158 | 20 |
| Etoxazole | 0.01 | 0.03 | 12.62 | 140 | Positive | 360.1 | 141 | 30 | 304 | 18 |
| Amitraz | 0.06 | 0.18 | 12.79 | 104 | Positive | 294.2 | 163.1 | 15 | 122 | 30 |
| Spiromesifen | 0.1 | 0.3 | 12.90 | 76 | Positive | 371.2 | 273.1 | 10 | 255.1 | 23 |
| Avermectin B1a | 0.3 | 0.9 | 12.91 | 176 | Positive | 890.3 | 305.2 | 24 | 567.3 | 13 |

**Table S2.** Pesticide properties of the active ingredients detected in soil samples. DT50 = half-life time in soil (days), Sw = solubility in water at 20 °C (mg/L), LogP = octanol-water partition coefficient at pH 7 and 20°C, Vp = vapor pressure at 20 °C (mPa), GUS = GUS index, Koc = organic carbon-water partition coefficient (ml/g).

| Active ingredient | DT50 | Sw | LogP | Vp | Koc | GUS |
|---|---|---|---|---|---|---|
| 4-Hydroxy-chlorothalonil | 3.53 | 0.81 | 2.94 | 0.076 | 2632 | 1.12 |
| Acetamiprid | 1.6 | 2950 | 0.8 | 0.000173 | 200 | 0.94 |
| Ametryn | 37 | 200 | 2.63 | 0.365 | 316 | 0.46 |
| Atrazine | 75 | 35 | 2.7 | 0.039 | 100 | 2.57 |
| Azoxystrobin | 78 | 6.7 | 2.5 | 0.00000011 | 589 | 3.1 |
| Boscalid | 484.4 | 4.6 | 2.96 | 0.00072 | 9500 [a] | 2.68 |
| Carbaryl | 16 | 9.1 | 2.36 | 0.0416 | 300 | 2.02 |
| Carbofuran | 29 | 322 | 1.8 | 0.08 | 65.15 [a] | 2.36 |
| Chlorantraniliprole | 597 | 0.88 | 2.86 | 6.3E-09 | 362 | 3.51 |
| Chloridazon | 31 | 422 | 1.19 | 0.000001 | 120 | 2.16 |
| Chloroxuron | 36 | 3.7 | 3.4 | 0.00023 | 2820 | 0.98 |
| Clomazone | 22.6 | 1212 | 2.58 | 27 | 300 | 2.72 |
| Clothianidin | 545 | 340 | 0.905 | 2.8E-08 | 123 | 3.74 |
| Cyantraniliprole | 34.4 | 14.2 | 2.02 | 5.13E-12 | 241 | 2.59 |
| Cyprodinil | 37 | 13 | 4 | 0.51 | 2829.5 [a] | 1.06 |
| Difenoconazole | 130 | 15 | 4.36 | 0.0000333 | 5467 [a] | 0.83 |
| Dimethenamid | 13 | 1200 | 2.2 | 0.37 | 140 [a] | 2.41 |
| Dimethomorph | 72.7 | 28.95 | 2.68 | 0.00097 | 5690 [a] | 2.26 |
| Diuron | 146.6 | 35.6 | 2.87 | 0.00115 | 680 | 2.65 |
| Fenbuconazole | 60 | 2.47 | 3.79 | 0.00034 | 2185 [b] | 0.63 |
| Fluazinam | 124 | 0.135 | 4.87 | 0.0172 | 16430 | 1 |
| Fludioxonil | 164 | 1.8 | 4.12 | 0.00039 | 145600 | -1.35 |
| Fluopicolide | 271 | 2.8 | 2.9 | 0.000303 | 190.65 [a] | 3.2 |
| Fluopyram | 309 | 16 | 3.3 | 0.0012 | 363 [b] | 3.23 |
| Flusilazole | 300 | 41.9 | 3.87 | 0.0387 | 1664 | 1.54 |
| Fluxapyroxad | 183 | 3.44 | 3.13 | 0.0000027 | 950 [b] | 2.57 |
| Griseofulvin | NA | 8.64 | 2.36 | 0.00128 | NA | NA |
| Imidacloprid | 191 | 610 | 0.57 | 0.0000004 | 478 [a] | 3.69 |
| Kresoxim-methyl | 16 | 2 | 3.4 | 0.0023 | 1700 [a] | 0 |
| Metalaxyl | 36 | 84000 | 1.75 | 0.75 | 162 | 2.06 |
| Metconazole | 142.2 | 30.4 | 3.85 | 0.000021 | 1874.5 [b] | 2.03 |
| Methoxyfenozide | 456 | 3.3 | 3.72 | 0.0133 | 402 | 3 |
| Metolachlor | 90 | 530 | 3.4 | 1.7 | 120 | 2.36 |
| Metrafenone | 200.9 | 0.492 | 4.3 | 0.153 | 7061 | 0.91 |
| Myclobutanil | 560 | 132 | 2.89 | 0.198 | 950 [a] | 1.99 |
| Napropamide | 70 | 74 | 3.3 | 0.022 | 839 | 1.96 |
| Penthiopyrad | 121.5 | 1.375 | 4.62 | 0.000643 | 804 | 1.08 |
| Phosmet | 3.2 | 15.2 | 2.8 | 0.065 | 3534 | 0.48 |
| Piperonyl butoxide | 13 | 14.3 | 4.75 | 0.02 | 89125 | -1.06 |
| Prometon | 500 | 620 | 2.91 | 0.306 | 43.2 | 6.31 |

| | | | | | | |
|---|---|---|---|---|---|---|
| Prometryn | 41 | 33 | 3.34 | 0.13 | 400 | 0.59 |
| Propazine | 131 | 8.6 | 3.95 | 0.004 | 154 | 3.86 |
| Propiconazole | 71.8 | 150 | 3.72 | 0.056 | 1086 | 1.58 |
| Pyraclostrobin | 41.9 | 1.9 | 3.99 | 0.000026 | 9304 | 0.05 |
| Pyrimethanil | 50.9 | 110 | 2.84 | 1.1 [a] | 835 | 2.17 |
| Spinetoram | 16.1 | 29 | 4.2 | 0.057 | 22836 | 0.27 |
| Tebufenozide | 400 | 0.83 | 4.25 | 0.000156 | 35000 [a] | 1.72 |
| Tebuthiuron | 400 | 2500 | 1.79 | 0.27 | 80 | 5.36 |
| Thiabendazole | 500 | 30 | 2.39 | 0.00053 | 3983 | 1.94 |
| Thiacloprid | 0.88 | 184 | 1.26 | 0.0000003 | 1100 [a] | 1.1 |
| Thiamethoxam | 50 | 4100 | -0.13 | 0.0000066 | 56.2 | 3.58 |
| Thiophanate-methyl | 0.5 | 18.5 | 1.4 | 0.009 | 330 [a] | 0.5 |
| Triadimenol | 250 | 72 | 3.18 | 0.0005 | 750 | 2.44 |
| Trifloxystrobin | 0.34 | 0.61 | 4.5 | 0.0034 | 3325.75 [a] | 0.15 |

All pesticide properties were obtained from the Pesticide Properties Database of the University of Hertfordshire (Lewis et al. 2016), except the values indicated by: [a] PubChem (Kim et al. 2020), and [b] US EPA Pesticide Fact Sheets (US EPA 2021).

**Table S3**. Frequency of detection (%), maximum, minimum, mean, and median concentrations (ng/g) of the pesticide residues detected in agricultural soils sampled at potential hibernation sites for bumblebee queens in Ontario, Canada. n = number of soil samples with detections (≥LOD).

| | Use | Apple orchards (n=20) | | | | Diversified farms (n=46) | | | | All samples (n=66) | | | | |
|---|---|---|---|---|---|---|---|---|---|---|---|---|---|---|
| | | Max | Mean (SD) | Median | % (n) | Max | Mean (SD) | Median | % (n) | Min* | Max | Mean (SD) | Median | % (n) |
| 4-Hydroxy-chlorothalonil | F | 151.79 | 47.39 (70.42) | 17.38 | 20.0 (4) | NA | NA | NA | *0.0 (0)* | *<LOQ* | 151.79 | 47.39 (70.42) | 17.38 | 6.1 (4) |
| Acetamiprid | I | 0.53 | 0.33 (0.29) | 0.33 | 10.0 (2) | NA | NA | NA | 0.0 (0) | <LOQ | 0.53 | 0.33 (0.29) | 0.33 | 3.0 (2) |
| Ametryn | H | <LOQ | 0.06 | 0.06 | 5.0 (1) | NA | NA | NA | 0.0 (0) | <LOQ | <LOQ | 0.06 | 0.06 | 1.5 (1) |
| Atrazine | H | 12.41 | 1.20 (3.39) | 0.15 | 65.0 (13) | 5.01 | 0.60 (1.13) | 0.18 | 87.0 (40) | <LOQ | 12.41 | 0.75 (1.92) | 0.18 | 80.3 (53) |
| Azoxystrobin | F | 8.53 | 0.97 (2.52) | 0.08 | 55.0 (11) | 0.22 | 0.08 (0.06) | 0.06 | 15.2 (7) | <LOQ | 8.53 | 0.62 (1.98) | 0.06 | 27.3 (18) |
| Boscalid | F | >1000.00 | 184.73 (346.19) | 20.1 | 70.0 (14) | 102.00 | 16.67 (25.89) | 4.80 | 32.6 (15) | <LOQ | >1000.00 | 97.80 (251.56) | 13.74 | 43.9 (29) |
| Carbaryl | I | 1.68 | 0.69 (0.59) | 0.30 | 45.0 (9) | 0.17 | 0.16 (0.01) | 0.15 | 6.5 (3) | <LOQ | 1.68 | 0.56 (0.56) | 0.28 | 18.2 (12) |
| Carbofuran | I | 0.11 | 0.11 | 0.11 | 5.0 (1) | 0.42 | 0.14 (0.14) | 0.07 | 13.0 (6) | <LOQ | 0.42 | 0.13 (0.13) | 0.08 | 10.6 (7) |
| Chlorantraniliprole | I | 171.31 | 24.99 (43.97) | 3.27 | 100.0 (20) | 51.43 | 2.62 (9.65) | 0.30 | 60.9 (28) | <LOQ | 171.31 | 11.94 (30.97) | 0.77 | 72.7 (48) |
| Chloridazon | H | 3.84 | 3.84 | 3.84 | 5.0 (1) | NA | NA | NA | 0.0 (0) | 3.84 | 3.84 | 3.84 | 3.84 | 1.5 (1) |
| Chloroxuron | H | NA | NA | NA | 0.0 (0) | 0.39 | 0.39 | 0.39 | 2.2 (1) | 0.39 | 0.39 | 0.39 | 0.39 | 1.5 (1) |
| Clomazone | H | 4.51 | 4.51 | 4.51 | 5.0 (1) | 2.73 | 1.10 (1.42) | 0.38 | 6.5 (3) | <LOQ | 4.51 | 1.95 (2.06) | 1.55 | 6.1 (4) |
| Clothianidin | I | 1.07 | 0.72 (0.20) | 0.60 | 30.0 (6) | 1.04 | 0.65 (0.15) | 0.60 | 19.6 (9) | <LOQ | 1.07 | 0.68 (0.17) | 0.60 | 22.7 (15) |
| Cyantraniliprole | I | 148.82 | 18.17 (41.42) | 0.74 | 65.0 (13) | <LOQ | 0.30 | 0.30 | 2.2 (1) | <LOQ | 148.82 | 16.89 (40.08) | 0.72 | 21.2 (14) |
| Cyprodinil | F | 4.87 | 3.21 (2.35) | 3.21 | 10.0 (2) | 0.30 | 0.23 (0.13) | 0.30 | 6.5 (3) | 0.08 | 4.87 | 1.42 (2.01) | 0.30 | 7.6 (5) |
| Difenoconazole | F | 25.89 | 7.51 (10.50) | 4.70 | 25.0 (5) | NA | NA | NA | 0.0 (0) | 0.73 | 25.89 | 7.51 (10.50) | 4.70 | 7.6 (5) |
| Dimethenamid | H | NA | NA | NA | 0.0 (0) | 0.22 | 0.22 | 0.22 | 2.2 (1) | 0.22 | 0.22 | 0.22 | 0.22 | 1.5 (1) |

| | | | | | | | | | | | | | | |
|---|---|---|---|---|---|---|---|---|---|---|---|---|---|---|
| Dimethomorph | F | 0.28 | 0.23 (0.05) | 0.24 | 15.0 (3) | NA | NA | NA | 0.0 (0) | <LOQ | 0.28 | 0.23 (0.05) | 0.24 | 7.6 (5) |
| Diuron | H | <LOQ | 0.90 | 0.90 | 10.0 (2) | NA | NA | NA | 0.0 (0) | <LOQ | <LOQ | 0.9 | 0.9 | 3.0 (2) |
| Fenbuconazole | F | 10.57 | 6.30 (6.04) | 6.30 | 10.0 (2) | NA | NA | NA | 0.0 (0) | 2.03 | 10.57 | 6.30 (6.04) | 6.30 | 3.0 (2) |
| Fluazinam | F | <LOQ | 0.30 | 0.30 | 5.0 (1) | NA | NA | NA | 0.0 (0) | <LOQ | <LOQ | 0.30 | 0.30 | 1.5 (1) |
| Fludioxonil | F | 44.58 | 44.58 | 44.58 | 5.0 (1) | 1.16 | 1.16 | 1.16 | 2.2 (1) | 1.16 | 44.58 | 22.87 (30.70) | 22.87 | 3.0 (2) |
| Fluopicolide | F | 0.26 | 0.22 (0.06) | 0.22 | 10.0 (2) | NA | NA | NA | 0.0 (0) | <LOQ | 0.26 | 0.22 (0.06) | 0.22 | 3.0 (2) |
| Fluopyram | F | 93.42 | 6.97 (21.23) | 0.27 | 95.0 (19) | 0.14 | 0.09 (0.05) | 0.06 | 6.5 (3) | <LOQ | 93.42 | 6.03 (19.80) | 0.21 | 33.3 (22) |
| Flusilazole | F | 1.10 | 0.49 (0.43) | 0.18 | 40.0 (8) | NA | NA | NA | 0.0 (0) | <LOQ | 1.10 | 0.49 (0.43) | 0.18 | 12.1 (8) |
| Fluxapyroxad | F | 1.33 | 0.59 (0.55) | 0.23 | 25.0 (5) | NA | NA | NA | 0.0 (0) | <LOQ | 1.33 | 0.59 (0.55) | 0.23 | 7.6 (5) |
| Griseofulvin | F | <LOQ | 0.30 | 0.30 | 5.0 (1) | 0.30 | 0.30 | 0.30 | 2.2 (1) | <LOQ | 0.30 | 0.30 | 0.30 | 3.0 (2) |
| Imidacloprid | I | 0.41 | 0.35 (0.05) | 0.34 | 15.0 (3) | 3.97 | 0.71 (1.09) | 0.30 | 23.9 (11) | <LOQ | 3.97 | 0.63 (0.97) | 0.31 | 21.2 (14) |
| Kresoxim-methyl | F | 1.08 | 1.08 | 1.08 | 5.0 (1) | NA | NA | NA | 0.0 (0) | 1.08 | 1.08 | 1.08 | 1.08 | 1.50 (1) |
| Metalaxyl | F | 0.25 | 0.16 (0.13) | 0.16 | 10.0 (2) | NA | NA | NA | 0.0 (0) | <LOQ | 0.25 | 0.16 (0.13) | 0.16 | 3.0 (2) |
| Metconazole | F | 42.98 | 42.98 | 42.98 | 5.0 (1) | NA | NA | NA | 0.0 (0) | 42.98 | 42.98 | 42.98 | 42.98 | 1.5 (1) |
| Methoxyfenozide | I | 23.89 | 4.78 (8.85) | 0.59 | 35.0 (7) | 0.73 | 0.44 (0.41) | 0.44 | 4.3 (2) | <LOQ | 23.89 | 3.82 (7.90) | 0.59 | 13.6 (9) |
| Metolachlor | H | 173.09 | 15.59 (46.54) | 0.38 | 70.0 (14) | 17.77 | 1.88 (3.56) | 0.83 | 60.9 (28) | <LOQ | 173.09 | 6.45 (27.17) | 0.58 | 63.6 (42) |
| Metrafenone | F | <LOQ | 0.18 | 0.18 | 5.0 (1) | NA | NA | NA | 0.0 (0) | <LOQ | <LOQ | 0.18 | 0.18 | 1.5 (1) |
| Myclobutanil | F | 1.65 | 0.47 (0.58) | 0.21 | 30.0 (6) | NA | NA | NA | 0.0 (0) | <LOQ | 1.65 | 0.47 (0.58) | 0.21 | 9.1 (6) |
| Napropamide | H | <LOQ | 0.06 | 0.06 | 5.0 (1) | 13.92 | 1.69 (3.60) | 0.68 | 30.4 (14) | <LOQ | 13.92 | 1.58 (3.50) | 0.53 | 22.7 (15) |
| Penthiopyrad | F | 50.71 | 6.91 (17.72) | 0.18 | 40.0 (8) | NA | NA | NA | 0.0 (0) | 0.07 | 50.71 | 6.91 (17.73) | 0.18 | 12.1 (8) |
| Phosmet | I | 8.28 | 2.94 (2.99) | 1.50 | 25.0 (5) | NA | NA | NA | 0.0 (0) | <LOQ | 8.28 | 2.94 (2.99) | 1.50 | 7.6 (5) |

| | | | | | | | | | | | | | | |
|---|---|---|---|---|---|---|---|---|---|---|---|---|---|---|
| Piperonyl butoxide | S | NA | NA | NA | 0.0 (0) | 0.06 | 0.06 | 0.06 | 2.2 (1) | 0.06 | 0.06 | 0.06 | 0.06 | 1.5 (1) |
| Prometon | H | NA | NA | NA | 0.0 (0) | 0.37 | 0.37 | 0.37 | 2.2 (1) | 0.37 | 0.37 | 0.37 | 0.37 | 1.5 (1) |
| Prometryn | H | NA | NA | NA | 0.0 (0) | <LOQ | 0.06 | 0.06 | 2.2 (1) | <LOQ | <LOQ | 0.06 | 0.06 | 1.5 (1) |
| Propazine | H | 0.27 | 0.27 | 0.27 | 5.0 (1) | <LOQ | 0.06 | 0.06 | 6.5 (3) | <LOQ | 0.27 | 0.11 (0.11) | 0.06 | 6.1 (4) |
| Propiconazole | F | 11.58 | 6.80 (6.76) | 6.80 | 10.0 (2) | <LOQ | 0.06 | 0.06 | 2.2 (1) | <LOQ | 11.58 | 4.55 (6.16) | 2.02 | 4.5 (3) |
| Pyraclostrobin | F | 147.18 | 22.35 (45.13) | 2.00 | 70.0 (14) | 2.57 | 0.61 (0.80) | 0.26 | 21.7 (10) | <LOQ | 147.18 | 13.29 (35.65) | 0.81 | 36.4 (24) |
| Pyrimethanil | F | 8.51 | 1.88 (3.04) | 0.32 | 35.0 (7) | NA | NA | NA | 0.0 (0) | <LOQ | 8.51 | 1.88 (3.04) | 0.32 | 10.6 (7) |
| Spinetoram | I | 0.73 | 0.36 (0.32) | 0.18 | 15.0 (3) | 1.36 | 1.36 | 1.36 | 2.2 (1) | <LOQ | 1.36 | 0.61 (0.56) | 0.46 | 6.1 (4) |
| Tebufenozide | I | 13.98 | 4.75 (7.99) | 0.22 | 15.0 (3) | NA | NA | NA | 0.0 (0) | <LOQ | 13.98 | 4.83 (7.92) | 0.30 | 4.5 (3) |
| Tebuthiuron | H | 1.15 | 1.15 | 1.15 | 5.0 (1) | NA | NA | NA | 0.0 (0) | 1.15 | 1.15 | 1.15 | 1.15 | 1.5 (1) |
| Thiabendazole | F | <LOQ | 0.15 | 0.15 | 5.0 (1) | <LOQ | 0.15 | 0.15 | 4.3 (2) | <LOQ | <LOQ | 0.15 | 0.15 | 4.5 (3) |
| Thiacloprid | I | 5.67 | 0.89 (1.81) | 0.25 | 45.0 (9) | 0.19 | 0.19 | 0.19 | 2.2 (1) | <LOQ | 5.67 | 0.82 (1.73) | 0.22 | 15.1 (10) |
| Thiamethoxam | I | <LOQ | 0.18 | 0.18 | 5.0 (1) | <LOQ | 0.18 | 0.18 | 2.2 (1) | <LOQ | <LOQ | 0.18 | 0.18 | 3.0 (2) |
| Thiophanate-methyl | F | <LOQ | 0.06 | 0.06 | 5.0 (1) | NA | NA | NA | 0.0 (0) | <LOQ | <LOQ | 0.06 | 0.06 | 1.5 (1) |
| Triadimenol | F | 14.53 | 4.95 (8.30) | 0.18 | 15.0 (3) | NA | NA | NA | 0.0 (0) | 0.14 | 14.53 | 4.95 (8.30) | 0.18 | 4.5 (3) |
| Trifloxystrobin | F | 0.18 | 0.14 (0.05) | 0.16 | 15.0 (3) | <LOQ | 0.06 | 0.06 | 2.2 (1) | <LOQ | 0.18 | 0.13 (0.06) | 0.13 | 6.1 (4) |

Use: F = fungicide, H = herbicide, I = insecticide, S = pesticide synergist
* LOQ values are available in table S1
NA = not applicable

**Table S4**. Combinations of active ingredients detected in the 66 soil samples. n = number of soil samples presenting the respective pesticide combination.

| Number of active ingredients in soil (>LOD) | n |
|---|---|
| Diversified farms (n=46) | |
| 0 residue | 1 |
| 1 residue | 5 |
| Atrazine | 4 |
| Napropamide | 1 |
| 2 residues | 9 |
| Atrazine + chlorantraniliprole | 3 |
| Atrazine + clothianidin | 1 |
| Atrazine + metolachlor | 3 |
| Atrazine + prometon | 1 |
| Chlorantraniliprole + metolachlor | 1 |
| 3 residues | 8 |
| Atrazine + chlorantraniliprole + clothianidin | 2 |
| Atrazine + chlorantraniliprole + metolachlor | 3 |
| Atrazine + imidacloprid + propiconazole | 1 |
| Atrazine + metolachlor + propazine | 2 |
| 4 residues | 5 |
| Atrazine + carbaryl + clothianidin + metolachlor | 1 |
| Atrazine + carbofuran + chlorantraniliprole + metolachlor | 1 |
| Atrazine + chlorantraniliprole + boscalid + pyraclostrobin | 1 |
| Atrazine + chlorantraniliprole + clothianidin + metolachlor | 1 |
| Atrazine + imidacloprid + metolachlor + napropamide | 1 |
| 5 residues | 3 |
| Atrazine + clothianidin + imidacloprid + metolachlor + napropamide | 1 |
| Boscalid + chlorantraniliprole + griseofulvin + metolachlor + napropamide | 1 |
| Carbaryl + chlorantraniliprole + cyantraniliprole + fluopyram + thiacloprid | 1 |
| 6 residues | 5 |
| Atrazine + azoxystrobin + boscalid + carbaryl + carbofuran + chlorantraniliprole | 1 |
| Atrazine + azoxystrobin + chlorantraniliprole + clothianidin + metolachlor + thiamethoxam | 1 |
| Atrazine + azoxystrobin + chlorantraniliprole + dimethenamid + metolachlor + propazine | 1 |
| Atrazine + boscalid + chlorantraniliprole + cyprodinil + fluopyram + pyraclostrobin | 1 |
| Atrazine + boscalid + clomazone + imidacloprid + metolachlor + pyraclostrobin | 1 |

| | |
|---|---|
| 7 residues | 3 |
| Atrazine + boscalid + chlorantraniliprole + chloroxuron + imidacloprid + metolachlor + napropamide | 1 |
| Atrazine + boscalid + chlorantraniliprole + imidacloprid + metolachlor + napropamide + pyraclostrobin | 1 |
| Boscalid + chlorantraniliprole + cyprodinil + fludioxonil + metolachlor + napropamide + pyraclostrobin | 1 |
| 8 residues | 2 |
| Atrazine + azoxystrobin + boscalid + chlorantraniliprole + clothianidin + metolachlor + napropamide + pyraclostrobin | 1 |
| Atrazine + boscalid + carbofuran + chlorantraniliprole + imidacloprid + metolachlor + napropamide + pyraclostrobin | 1 |
| 9 residues | 2 |
| Atrazine + azoxystrobin + boscalid + chlorantraniliprole + clomazone + imidacloprid + metolachlor + napropamide + prometryn | 1 |
| Atrazine + boscalid + chlorantraniliprole + clothianidin + cyprodinil + metolachlor + napropamide + pyraclostrobin + spinetoram | 1 |
| 10 residues | 1 |
| Atrazine+ azoxystrobin + boscalid + carbofuran + chlorantraniliprole + imidacloprid + metolachlor + methoxyfenozide + napropamide + thiabendazole | 1 |
| 11 residues | 1 |
| Atrazine + azoxystrobin + boscalid + carbofuran + chlorantraniliprole + fluopyram + imidacloprid + methoxyfenozide + metolachlor + napropamide + pyraclostrobin | 1 |
| 12 residues | 1 |
| Atrazine + boscalid + carbofuran + chlorantraniliprole + clomazone + imidacloprid + metolachlor + napropamide + piperonyl butoxide + pyraclostrobin + thiabendzole + trifloxystrobin | 1 |
| Apple orchards (n=20) | |
| 5 residues | 1 |
| Atrazine + boscalid + chlorantraniliprole + fluopyram + penthiopyrad | 1 |
| 6 residues | 2 |
| Atrazine + azoxystrobin + carbaryl + chlorantraniliprole + cyantraniliprole + fluopyram | 1 |
| Boscalid + chlorantraniliprole + cyantraniliprole + fluopyram + metolachlor + thiacloprid | 1 |
| 7 residues | 2 |
| Atrazine + azoxystrobin + carbofuran + chlorantraniliprole + clothianidin + fluopyram + metolachlor | 1 |
| Atrazine + azoxystrobin + chlorantraniliprole + cyantraniliprole + fluopyram + imidacloprid + thiabendazole | 1 |
| 8 residues | 1 |
| Boscalid + chlorantraniliprole + cyantraniliprole + fluopyram + metolachlor + penthiopyrad + pyraclostrobin + thiacloprid | 1 |

| | | |
|---|---|---|
| 9 residues | | 3 |
| | Acetamiprid + ametryn + atrazine + carbaryl + chlorantraniliprole + fluopyram + fluxapyroxam + metolachlor + pyraclostrobin | 1 |
| | Atrazine + carbaryl + chlorantraniliprole + cyantraniliprole + fluopyram + imidacloprid + myclobutanil + pyrimethanil + thiacloprid | 1 |
| | Boscalid + carbaryl + chlorantraniliprole + clothianidin + cyantraniliprole + methoxyfenozide + metolachlor + pyraclostrobin + tebufenozide | 1 |
| 10 residues | | 1 |
| | Boscalid + chlorantraniliprole + cyantraniliprole + fluopyram + flusilazole + penthiopyrad + pyraclostrobin + pyrimethanil + spinetoram + thiacloprid | 1 |
| 11 residues | | 2 |
| | 4-hydroxy-chlorothalonil + atrazine + azoxystrobin + chlorantraniliprole + clothianidin + dimethomorph + fluopyram + metalaxyl + metolachlor + myclobutanil + pyraclostrobin | 1 |
| | Boscalid + carbaryl + chlorantraniliprole + cyantraniliprole + fluopyram + flusilazole + penthiopyrad + pyraclostrobin + pyrimethanil + spinetoram + thiacloprid | 1 |
| 13 residues | | 2 |
| | Atrazine + azoxystrobin + boscalid + chlorantraniliprole + dimethomorph + fluopicolide + fluopyram + flusilazole + methoxyfenozide + metolachlor + myclobutanil + pyraclostrobin + pyrimethanil | 1 |
| | Atrazine + azoxystrobin + boscalid + chlorantraniliprole + dimethomorph + fluopyram + fluxapyroxam + methoxyfenozide + metolachlor + myclobutanil + phosmet + pyraclostrobin + thiophanate-methyl | 1 |
| 14 residues | | 1 |
| | Atrazine + azoxystrobin + boscalid + carbaryl + clorantraniliprole + clothianidin + fluopyram + flusilazole + griseofulvin + metalaxyl + metolachlor + myclobutanil + pyraclostrobin + pyrimethanil | 1 |
| 19 residues | | 3 |
| | 4-hydroxy-chlorothalonil + acetamiprid + atrazine + azoxystrobin + boscalid + chlorantraniliprole + clothianidin + cyantraniliprole + difenoconazole + fenbuconazole + fluopyram + fluxapyroxide + imidacloprid + metolachlor + penthiopyrad + propiconazole + pyraclostrobin + thiacloprid + trifloxystobin | 1 |
| | Atrazine + azoxystrobin + boscalid + chlorantraniliprole + cyantraniliprole + cyprodinil + difenoconazole + fluopyram + flusilazole + krezoxym-methyl + methoxyfenozide + metolachlor + penthiopyrad + phosmet + pyrimethanil + pyraclostrobin + thiacloprid + triadimenol + trifloxystrobin | 1 |
| | Azoxystrobin + boscalid + carbaryl + chlorantraniliprole + cyantraniliprole + difenoconazole + diuron + fluazinam + fluopyram + flusilazole + methoxyfenozide + metolachlor + myclobutanil + napropamide + penthiopyrad + pyraclostrobin + tebuconazole + thiacloprid + triadimenol | 1 |
| 20 residues | | 1 |
| | Boscalid + carbaryl + chlorantraniliprole + cyantraniliprole + cyprodinil + difenoconazole + fludiaxonil + fluopyram + flusilazole + fluxapyroxam + metconazole + methoxyfenozide + metolachlor + metrafenone + phosmet | 1 |

| | | |
|---|---|---|
| | + propiconazole + pyraclostrobin + spinetoram + tebuconazole + triadimenol | |
| 29 residues | | 1 |
| | 4-hydroxy-chlorothalonil + acetamiprid + atrazine + azoxystrobin + boscalid + carbaryl + chlorantraniliprole + chloridazon + clomazone + clothianidin + cyantraniliprole + difenoconazole + diuron + fenbuconazole + fluopicolide + fluopyram + flusilazole + fluxapyroxam + methoxyfenozide + metolachlor + penthiopyrad + phosmet + propazine + pyraclostrobin + pyrimethanil + tebuthiuron + thiacloprid + thiamethoxam + trifloxystrobin | 1 |

**Table S5**. Spearman's correlation coefficients between soil properties and the total pesticide concentration in soil, the number of pesticide active ingredients per sample, or the concentration of the most common pesticide residues in soil (coefficients were calculated only when there was a minimum of five soil samples in which the AI was quantified; NA = not available). AO = apple orchards; DF = diversified farms; All = all soil samples. Positive values correspond to positive correlations and negative values to negative correlations. Significant correlations ($p < 0.05$) are marked in bold.

| | Type of cropping system | % clay | % silt | Organic matter content | soil pH | Soil humidity | Slope inclination | Distance to the nearest field |
|---|---|---|---|---|---|---|---|---|
| Total pesticide concentration | AO | -0.11 | 0.05 | -0.29 | 0.18 | -0.20 | -0.18 | **-0.49** |
| | DF | -0.16 | -0.10 | -0.20 | **-0.49** | -0.19 | -0.16 | -0.17 |
| | All | 0.12 | 0.22 | **-0.27** | -0.24 | -0.05 | **-0.25** | -0.22 |
| Number of pesticide active ingredients | AO | -0.38 | -0.13 | -0.24 | 0.07 | -0.40 | -0.09 | -0.32 |
| | DF | -0.12 | -0.08 | -0.22 | **-0.47** | -0.14 | -0.05 | -0.06 |
| | All | 0.16 | **0.28** | **-0.26** | -0.23 | 0.02 | -0.18 | -0.09 |
| Atrazine concentration | AO | -0.26 | -0.22 | -0.31 | 0.15 | -0.26 | 0.24 | -0.15 |
| | DF | -0.04 | 0.03 | 0.05 | 0.07 | -0.13 | -0.27 | -0.21 |
| | All | -0.08 | -0.03 | -0.02 | 0.08 | -0.17 | -0.14 | -0.19 |
| Azoxystrobin concentration | AO | -0.56 | -0.38 | -0.31 | 0.53 | 0.08 | 0.24 | -0.26 |
| | DF | -0.41 | -0.41 | -0.41 | -0.20 | 0.00 | 0.10 | -0.10 |
| | All | -0.23 | -0.05 | -0.44 | 0.38 | -0.02 | 0.12 | -0.20 |
| Boscalid concentration | AO | 0.13 | 0.10 | -0.35 | -0.11 | -0.12 | -0.18 | -0.50 |
| | DF | -0.26 | -0.42 | -0.40 | 0.01 | -0.47 | **-0.53** | -0.38 |
| | All | 0.02 | -0.02 | **-0.37** | 0.03 | -0.20 | -0.34 | **-0.43** |
| Chlorantraniliprole concentration | AO | 0.03 | 0.039 | -0.26 | 0.19 | 0.04 | -0.15 | -0.30 |
| | DF | -0.03 | -0.05 | 0.07 | -0.26 | 0.10 | 0.05 | 0.09 |
| | All | **0.30** | **0.36** | -0.15 | -0.02 | 0.18 | -0.15 | -0.05 |
| Clothianidin concentration | AO | 0.67 | -0.36 | **-0.86** | 0.51 | -0.01 | 0.51 | -0.43 |
| | DF | 0.56 | 0.55 | -0.55 | 0.00 | -0.41 | 0.14 | -0.41 |
| | All | **0.66** | 0.35 | **-0.70** | 0.24 | -0.30 | 0.26 | -0.40 |
| Cyantraniliprole concentration | AO | 0.45 | 0.37 | -0.13 | 0.09 | 0.01 | -0.41 | **-0.57** |
| | DF | NA | NA | NA | NA | NA | NA | NA |
| | All | **0.53** | 0.46 | -0.22 | 0.04 | 0.17 | -0.46 | **-0.54** |
| Fluopyram concentration | AO | 0.16 | 0.27 | -0.02 | **0.48** | 0.15 | 0.29 | 0.21 |
| | DF | NA | NA | NA | NA | NA | NA | NA |
| | All | 0.38 | **0.49** | -0.07 | **0.52** | 0.27 | 0.21 | 0.17 |
| Flusilazole concentration | AO | 0.58 | 0.34 | -0.22 | 0.33 | 0.16 | 0.26 | 0.00 |
| | DF | NA | NA | NA | NA | NA | NA | NA |
| | All | 0.58 | 0.34 | -0.22 | 0.33 | 0.16 | 0.26 | 0.00 |
| Imidacloprid concentration | AO | NA | NA | NA | NA | 0.50 | 0.50 | NA |
| | DF | 0.20 | 0.20 | -0.05 | -0.49 | -0.28 | -0.30 | -0.43 |
| | All | 0.35 | 0.34 | -0.04 | -0.45 | -0.14 | -0.37 | -0.49 |
| Metolachlor concentration | AO | 0.35 | 0.24 | -0.43 | 0.07 | -0.3 | 0.14 | -0.13 |
| | DF | 0.19 | 0.08 | 0.13 | 0.07 | 0.20 | 0.06 | -0.10 |
| | All | 0.21 | -0.14 | 0.01 | 0.05 | 0.01 | 0.14 | 0.08 |

| | | | | | | | | |
|---|---|---|---|---|---|---|---|---|
| Napropamide concentration | AO | NA | NA | NA | NA | NA | NA | NA |
| | DF | -0.25 | -0.21 | 0.24 | 0.00 | 0.18 | -0.03 | 0.31 |
| | All | -0.32 | -0.28 | 0.27 | 0.00 | 0.12 | 0.05 | 0.42 |
| Penthiopyrad concentration | AO | -0.18 | 0.02 | -0.62 | 0.57 | -0.43 | 0.20 | -0.14 |
| | DF | NA | NA | NA | NA | NA | NA | NA |
| | All | -0.18 | 0.02 | -0.61 | 0.57 | -0.43 | 0.20 | -0.14 |
| Pyraclostrobin concentration | AO | 0.33 | -0.05 | -0.15 | 0.04 | -0.24 | 0.00 | -0.28 |
| | DF | -0.14 | 0.17 | 0.13 | -0.23 | 0.12 | 0.02 | 0.26 |
| | All | 0.37 | 0.36 | -0.16 | 0.09 | -0.04 | -0.16 | -0.20 |

**Figure S1.** Location of the farms where soil samples were collected in Ontario, Canada. Created from Google MyMaps (google.com/maps).

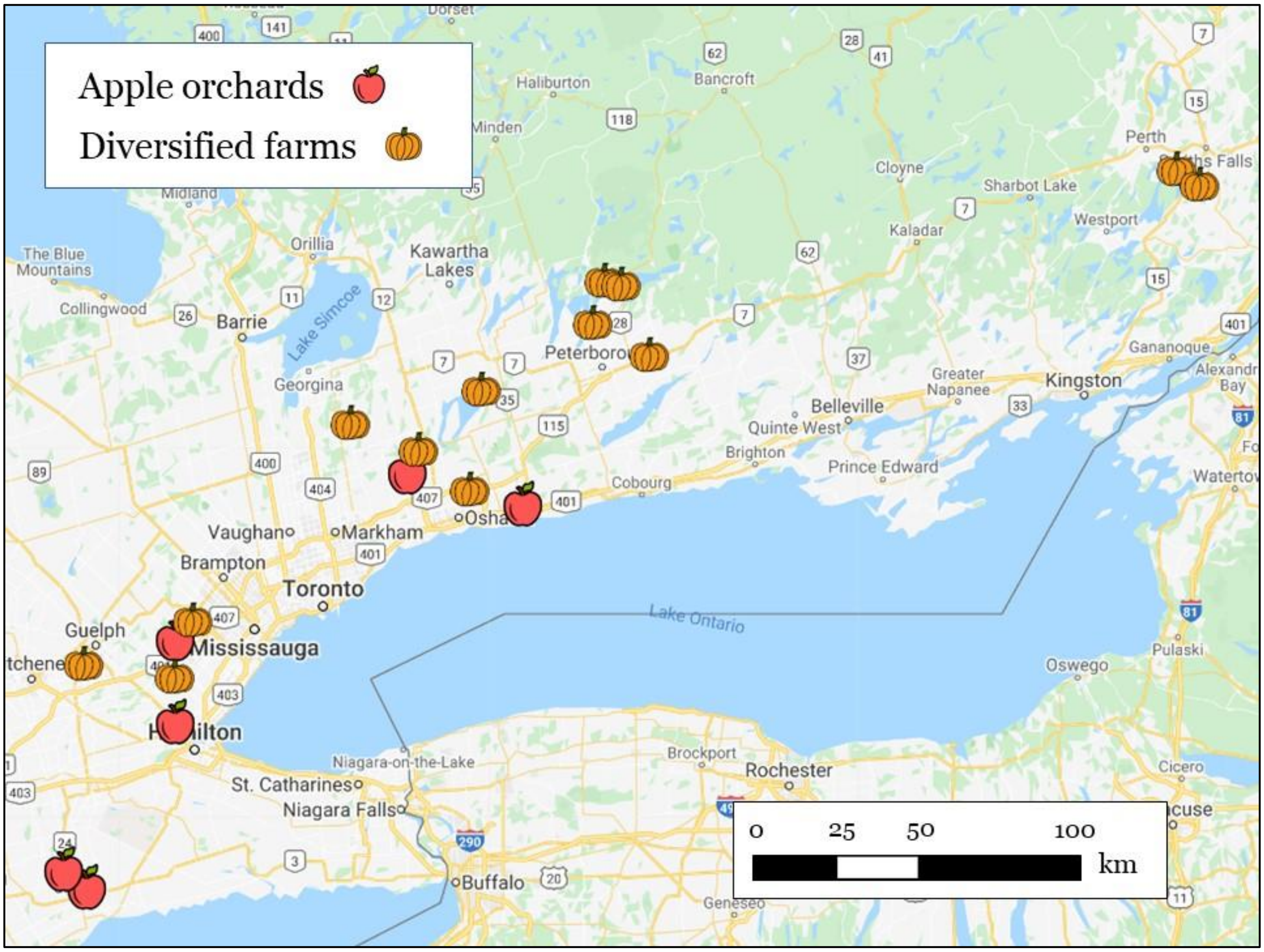
